# Unveiling the crucial role of betaine: Modulation of GABA homeostasis via SLC6A1 transporter (GAT1)

**DOI:** 10.1101/2024.04.30.591809

**Authors:** Manan Bhatt, Erika Lazzarin, Ana Sofia Alberto-Silva, Guido Domingo, Rocco Zerlotti, Ralph Gradisch, Andre Bazzone, Harald H. Sitte, Thomas Stockner, Elena Bossi

## Abstract

Betaine is an endogenous osmolyte that exhibits therapeutic potential by mitigating various neurological disorders. However, the underlying cellular and molecular mechanisms responsible for its neuroprotective effects remain puzzling.

In this study, we describe a possible mechanism behind the positive impact of betaine in preserving neurons from excitotoxicity. Using electrophysiology, mass spectroscopy, radiolabelled cellular assay, and molecular dynamics simulation we demonstrate that betaine at mM concentration acts as a slow substrate of GAT1 (*slc6a1*), the predominant GABA transporter in the central nervous system. Intriguingly, when betaine is present at low concentration (0.01-3 mM) with GABA (at concentration <K_0.5_), it blocks the GABA reuptake. This GAT1 modulation occurs through the temporal inhibition of the transporter, i.e., the prolonged occupancy by betaine impedes the rapid transition of the transporter to the inward conformation. The temporal inhibition results in a crucial regulatory mechanism contributing to the maintenance of GABA homeostasis, preserving neurons from excitotoxicity.

## Introduction

Betaine (N, N, N trimethylglycine) is a small molecule, found in diverse organisms from bacteria and plants to animals. It is a natural, stable, and nontoxic substance that can be produced endogenously in mitochondria by the oxidation of choline [1] and exogenously absorbed as a dietary nutrient from the betaine-rich food sources like beetroot, spinach, and different seafoods [2, 3]. In mammals, betaine is known to play two primary physiological roles: an osmolyte that protects the cells against osmotic pressure through osmoregulation, and an active methyl donor that enables the conversion of the toxic metabolite homocysteine into methionine [1]. In humans, betaine is mainly found in the liver, kidney, and brain. However, the presence and role of betaine in the brain is poorly understood and remains debated [4].

There has been substantial reporting on the beneficial effects of betaine supplementation in neurodegenerative and neuropsychiatric disorders like Alzheimer’s [5–7], Parkinson’s [8], schizophrenia [9], anhedonia [10], and others [4, 11, 12]. This reported evidence implies a role of betaine with therapeutic potential in the central nervous system (CNS) [4, 13]. While the evidence for the effects of betaine on neuronal disorders is strong, the cellular and molecular mechanisms involved in its membrane translocation remain unclear.

The solute carrier (SLC) membrane transporters regulate essential physiological functions like nutrient uptake, waste removal, and ion transport. This second-largest family of membrane proteins has much relevance to pharmacology as drug targets or mediators [14]. Betaine can enter the brain by crossing the blood-brain barrier (BBB) using betaine/γ-aminobutyric acid (GABA) transporter 1 (BGT-1, *slc6a12*) [15, 16]. However, the low expression of BGT-1 in the CNS raises questions over its effectiveness in betaine transport into the neurons [17]. The same is the case for another transporter called sodium-dependent amino acid transporter 2 (SNAT2, *slc38a2*), that transports betaine and is also expressed at low level in the CNS [18]. Interestingly, it has been published that betaine might interact with the GABAergic pathway beyond BGT-1 and play a role as a signalling ligand in the CNS [19–21]. To investigate the potential modulatory role of betaine, we studied its interaction with GABA transporter 1 (GAT1, *slc6a1*) [22]. The expression of GAT1 is mainly neuronal, predominantly in the adult frontal cortex, and among all GATs it is the most expressed GAT in the CNS. About 80% of GABA from the synaptic cleft is taken up into presynaptic neurons through GAT1 [23]. The modulation of the GABAergic activity via regulation of the GABA homeostasis sustains the excitatory/inhibitory (E/I) balance in the CNS, which is a critical factor in protecting the brain against the excitotoxicity and consequently reducing the risk of neurological and neuropsychiatric diseases [24].

The results reported here unveil a new role of betaine in the E/I balance by showing its important action of GAT1 in regulating GABA homeostasis. Here, we demonstrate a new relationship between GABA and betaine by integrating data from electrophysiology: two-electrode voltage clamp (TEVC) and automated patch clamp (Patchliner™), measurements of radiolabelled efflux assays, liquid chromatography-mass spectroscopy (LC-MS/MS), and molecular dynamics simulations.

## Results

### Extracellular betaine perfusion on *Xenopus laevis* oocytes heterologously expressing rGAT1 induces inward transport currents

Fully grown *Xenopus laevis* oocytes have a highly efficient biosynthetic apparatus that is able to perform all the post-translational modifications needed for correct protein targeting and function. We use the micro-injection technique to gain the heterologous expression of the protein of interest in *Xenopus laevis* oocytes[25]. BGT-1 is reported to transport betaine and GABA in a concentration-dependent manner, with higher affinity for GABA than betaine[26, 27]. The perfusion of betaine (1, 3, 10, 30, 50 mM) on the oocytes expressing rat GAT1 (rGAT1) resulted in concentration-dependent inward transport currents, like canine BGT-1 (cBGT-1) (Figure 1 A-C). At the holding potential V_h_ = -60 mV, the kinetic analysis of these inward transport currents provided a half maximal transport coefficient K_0.5_ = 11.57 ± 1.78 mM, the maximal transport current I_max_ = -76.15 ±3.46 nA, and the transport efficiency I_max_/K_0.5_ = -6.58 ± 0.80 nA/mM (Table in Figure 1). As a negative control, the non-injected oocytes were perfused with the same concentrations of betaine. Oocytes expressing rGAT1 and the non-injected control oocytes were also exposed to glycine (from 1 to 50 mM), and in both cases no inward current was observed (Figure S1 A). Additionally, also in CHO cells overexpressing human GAT1 (hGAT1) we recorded a concentration-dependent betaine transport current using the automated whole-cell patch clamp technique (Patchliner™ from Nanion GmbH, Germany) (Figure S2).

**Figure 1:**
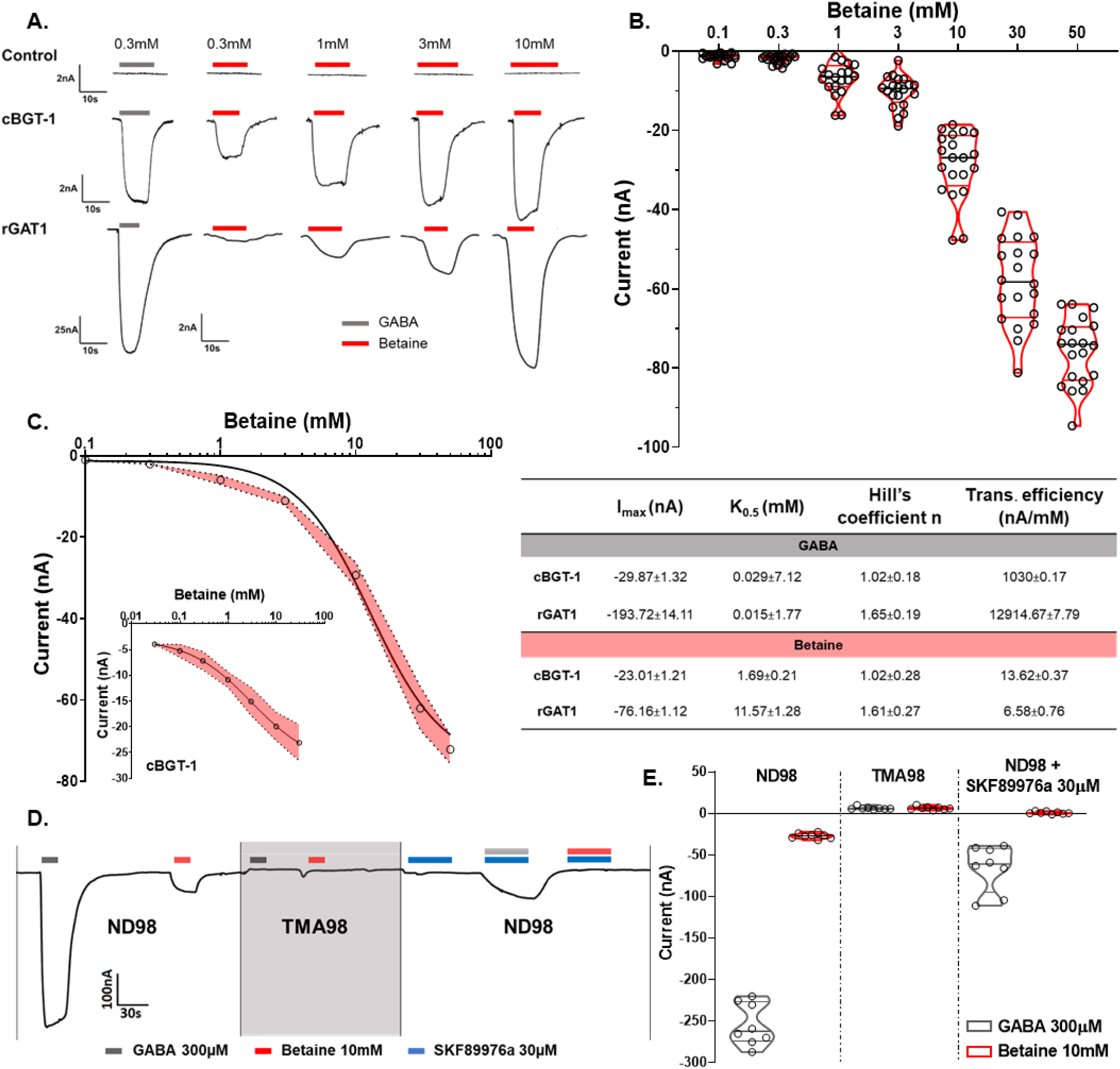
Betaine induces concentration- and sodium-dependent inward currents in Xenopus laevis oocytes expressing rGAT1, which can be blocked by SKF89976a. A. Representative traces of current recorded for GABA 300 µM and increasing betaine concentration (0.3-10 mM) in non-injected control oocytes, oocytes expressing cBGT-1, and rGAT1 (top to down). B. A violin scattered plot shows the concentration-dependent response of betaine (0.1-50 mM) in rGAT1. The current values are shown as mean ± SEM of 21/5 n (number of oocytes)/N (number of batches). C. The kinetic analysis of the betaine transport in rGAT1 yielded I_max_ = -76.16 ± 1.12 nA, K_0.5_ = 11.57 ± 1.28 mM, (see table). In the inset, for cBGT-1 the parameters were I_max_ = -23.01±1.21 nA, K_0.5_ = 1.69 ± 0.21 mM. All data were fitted using logistic fit model, with current values shown as mean ± SEM of 8/3 n/N. D. A representative trace of currents recorded for GABA 300 µM and betaine 10 mM in the presence of ND98, TMA98, and ND98+SKF89976a 30 µM. E. The histogram shows the mean values of the currents recorded as reported in panel D, the Na^+^ dependence, and the blocking effect of SKF89976a 30 µM on the inward induced currents by GABA 300 µM and betaine 10 mM. Current values are shown as mean ± SEM of 8/3 n/N. All recordings were performed at holding potential V_h_ = -60 mV.

### Betaine transport by rGAT1 is sodium-dependent and can be blocked by inhibitors of GAT1

Since GAT1 is a sodium-dependent symporter, meaning its transport activity requires a sodium gradient[28]. The replacement of sodium chloride (NaCl) 98 mM in the buffer solution (ND98) with trimethylammonium chloride (TMA-Cl) 98 mM resulted in the loss of the GABA- and betaine-induced inward transport currents in the oocyte heterologously expressing rGAT1 (Figure 1 D, E). We also tested the effects of SKF89976a 30 µM, a GAT1 inhibitor[29], on the current induced by GABA and betaine in the oocytes expressing rGAT1. The data showed strong inhibition of the transport current induced by GABA 300 µM and betaine 10 mM (Figure 1 D, E). Similar effects were also observed with other GAT1 inhibitors tiagabine and NO-711 (Figure S1 B), asserting further that the inward transport currents induced by betaine are mediated by rGAT1.

### The voltage-dependent transport of betaine by GAT1 shows that it is a slower substrate than GABA

GAT1 is a voltage-dependent transporter and the voltage-steps experiments on *X. laevis* oocytes expressing rGAT1 reveal pre-steady state, steady-state, and leak currents [30–32]. The voltage jump in the presence of betaine (1, 3, 10, 30, 50 mM) performed on the oocytes expressing rGAT1 elicited voltage-dependent transport currents in rGAT1, similar to that induced by GABA [33] (Figure 2 B).

**Figure 2:**
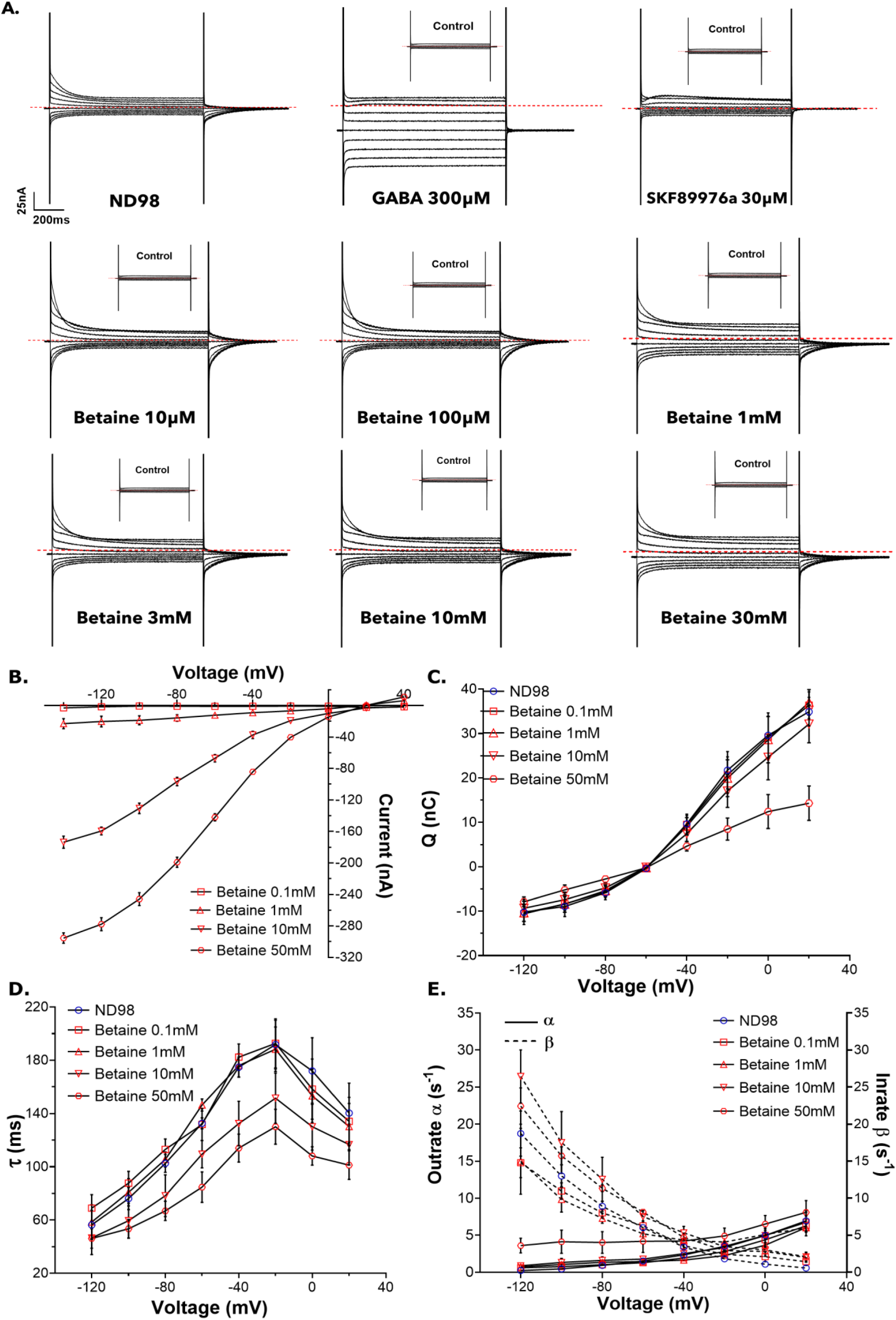
The pre-steady state analysis of betaine transport by rGAT1. The current response for each condition was collected by giving 0.8 s long squared pulse at –20 mV of from the holding potential of -60 mV, starting from –140 mV to +40 mV. **A.** The representative traces of the voltage-step response for ND98, GABA 300 µM, SKF89976a 30 µM, and different indicated betaine concentrations. The dashed red line indicates the holding current for the oocyte at the holding potential. **B.** The current and voltage (I-V) relationship from –140 mV to +40 mV. **C.** The total charge dislocation and voltage (Q-V) relationship. **D.** The decay time constant and voltage (τ-V) relationship. **E.** The relationships of unidirectional rate constants outrate (α) and inrate (β, shown as dashed line) with voltage. All the reported values were collected in the presence of ND98 alone and/or with betaine 0.1, 1, 10, 50 mM. In C-E the voltage levels tested were from –120 mV to +20 mV. For B-E, all values are shown as mean ± SEM of 3/1 n/N.

The pre-steady state (PSS) currents in GAT1 are due to the binding and unbinding of Na^+^ to the transporter inside the membrane electric field [34] and usually they disappear in the presence of a saturating concentration of substrate [31]. Hence expectedly, at saturating GABA concentration (300 µM), we observed the vanishing of PSS currents as they became too fast to be detected and only the steady state currents were detected. (Figure 2 A, top row). In the presence of SKF89976a (30 µM), the PSS and steady state both disappeared (Figure 2 A top row) as this compound blocks the access of Na^+^ to the GAT1 vestibule limiting the translocation of substrate and ions [35, 36]. The capacitive currents recorded in the presence of the inhibitors are the fast relaxation component due to the membrane capacitance of the oocytes. Interestingly, even in the presence of increasing betaine concentrations, we observed the persistence of a slow transient component. These relaxations are larger than expected (Figure 2 A, middle and last row) when compared to that at low GABA concentrations [37]. The PSS currents in the presence and absence of betaine were analysed after subtraction from the currents in the presence of SKF89976a 30 µM [25, 28] and the remainders were then integrated to obtain the dislocated charge (Q) and fitted with single exponential to determine the relaxation time constant (τ). The Q-V relationship reflected the charge moved into the membrane electric field at each tested transmembrane potential, whereas the τ-V relationship provided the rate of charge dislocation in the transporter vestibule. These data were further analysed to calculate outrate α and inrate β that indicated the rate of charge moving to and from the transporter cavity respectively (Figure 2 C, D, E) [37, 38]. All of the Q-V, τ-V, and rate constant analysis were done for betaine 0.1, 0.3, 1, 3, 10, 30, and 50 mM, in the figures the data for betaine 0.1, 1, 10, and 50 mM are reported (for the remaining concentrations Figure S3).

The Q-V relationship showed that low concentrations of betaine (0.1-1 mM) induce charge displacements similar to ND98 alone (Figure 2 C). With further increase in extracellular betaine (10 and 50 mM) the charge displacement decreased but less than expected and did not disappear like for saturating GABA [37]. The τ-V relationship provided a similar result, where the decay time constant decreased with the increase in betaine concentration (Figure 2 D). This indicates an accelerated transport rate in the presence of betaine but again differently from what is known for GABA [37]. Further analysis of β and α showed that at low betaine concentrations the charge entered the transporter cavities faster than it could leave (β>α), similar to the substrate free Na^+^- buffer solution. Interestingly, β in the presence of betaine 0.1 mM is lower than in the presence of Na^+^-buffer alone, in particular at voltages lower than -80 mV (Figure 2 E).

### Detection of GABA and betaine using LC-MS/MS in *X. laevis* oocytes expressing rGAT1

In this work, we developed a radiolabelled-free method using a tandem LC-MS/MS based approach to detect the substrates taken up by rGAT1 heterologously expressed in *Xenopus laevis* oocytes (Figure 3 A). To verify the sensitivity and linearity of the LC-MS/MS response to betaine and GABA, standard solutions were created, and a calibration curve was obtained (Figure S4). The detected retention times were 1’ 55” and 2’ 30” for GABA and betaine, respectively (Figure 3).

**Figure 3:**
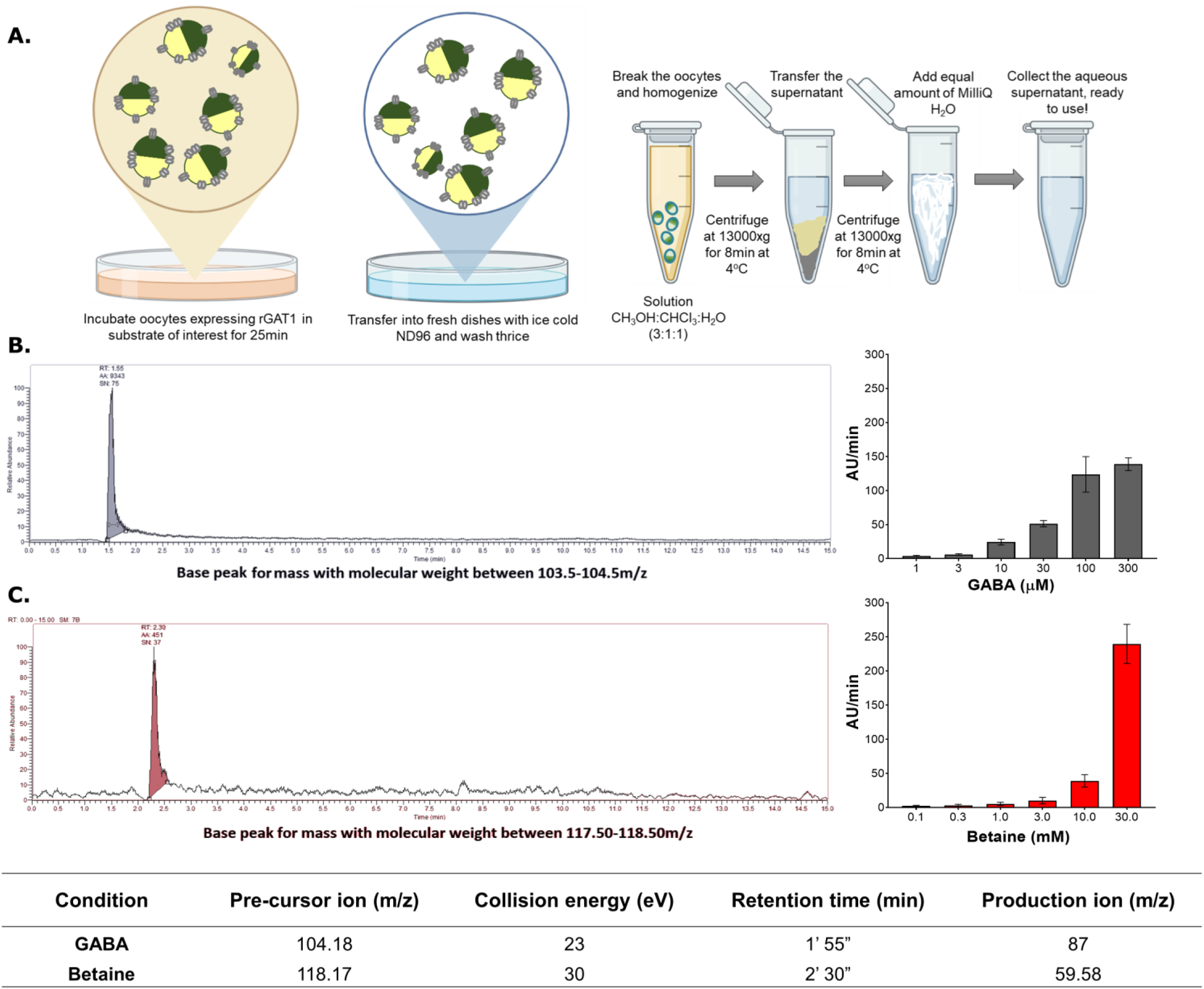
Detection of GABA and betaine transport by rGAT1 using the LC- MS/MS protocol on X. laevis oocytes. **A.** A cartoon of the protocol developed to extract the cytosol contents of the oocytes and detect the presence of the substrate of interest. **B.** A representative trace showing the presence of GABA in rGAT1 expressing oocytes incubated in GABA 1 mM. The histogram on the right shows a qualitative measurement of a concentration-dependent GABA uptake by rGAT1 expressing oocytes incubated in different GABA concentrations (1-300 µM). **C.** A representative trace showing the presence of betaine in rGAT1 expressing oocytes incubated in betaine 30 mM. The histogram on the right shows the qualitative measurement of a concentration-dependent betaine uptake by rGAT1 expressing oocytes incubated in different betaine concentrations (0.1-30 mM). All values are shown as arbitrary units per minute per oocyte ± SEM of 15/2 n/N. The table at the bottom shows the collision energy required to obtain a unique production ion and the retention time (in the 15-minute-long protocol) for the detection of the peak correlated to GABA and betaine.

The oocytes heterologously expressing rGAT1 were incubated in different concentrations of GABA (1, 3, 10, 30, 100, 300 µM) and betaine (0.1, 0.3, 1, 3, 10, 30 mM) for 25 minutes in groups of five oocytes per sample. For both GABA and betaine, a concentration-dependent uptake by rGAT1 was observed (Figure 3 B, C histogram on right). As a negative control, the non-injected oocytes were also incubated in the same concentrations of GABA and betaine, and their analysis showed absence of uptake.

### Both GABA and betaine induce efflux of [^3^H]-GABA in rGAT1 expressing HEK293 cells

We tested the effects of betaine on HEK293-rGAT1 cells pre-loaded with 0.01 µM [^3^H]- GABA at 37°C for 20 minutes. The experiment was initiated by replacing the pre- loading buffer with plain Kerbs Ringer HEPES buffer (KHB) and starting the experiment with the collection of baseline efflux values (Figure 4). Basal [^3^H]-GABA efflux was 0.12 ± 0.02% min^-1^. Addition of betaine induced a time-, concentration-, and rGAT1-dependent efflux of [^3^H]-GABA, in the presence and absence of the ionophore monensin (10 µM) (Figure 4 D, E). Monensin (mon) is a sodium ionophore that selectively collapses the Na^+^ and H^+^ gradients, reducing the electrochemical driving force for uptake and favouring the transporter efflux by an increase of sodium inside the cell [39]. As a positive control, pre-loaded HEK293-rGAT1 cells were exposed to GABA, and the results matched with the time-, concentration-, and rGAT1-dependent efflux (Figure 4 A, B), similar to our previous report [40]. For kinetic analysis, the drug-induced efflux was calculated as the mean efflux of the fraction where the value started plateauing divided by the length of the fraction (=2 min). The negative control was performed with tiagabine as the uptake inhibitor of rGAT1 (Figure 4 G). The fitting of the data using non-linear regression by the logistic fitting model provided the parameters as K_0.5,GABA_ = 36.28 ± 11.27 µM (with mon 10 µM K_0.5,GABA_ = 36.48 ± 2.57 µM) and K_0.5,betaine_ = 6.73 ± 2.21 mM (with mon 10 µM K_0.5,betaine_ = 7.71 ± 1.13 mM) (Figure 4 C, F).

**Figure 4:**
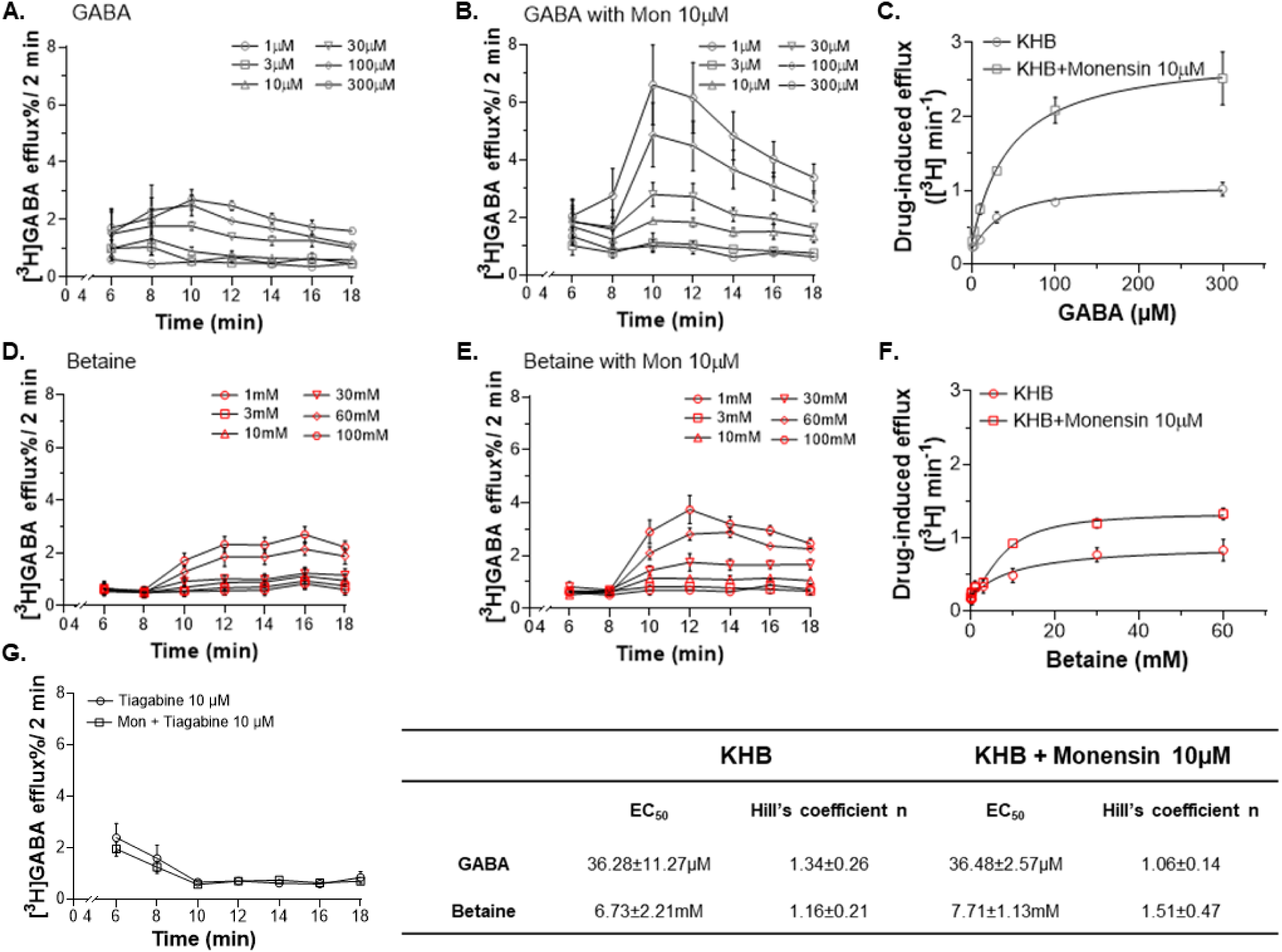
Betaine induces efflux of [^3^H]GABA in pre-loaded HEK293 cells overexpressing rGAT1. **A.** Time course of the efflux of [^3^H]GABA in the presence of increasing GABA concentrations (1-300 µM). **B.** Time course of the efflux of [^3^H]GABA in the presence of monensin 10 µM and different GABA concentrations (1-300 µM). **C.** The kinetic analysis of the [^3^H]GABA-induced efflux with and without monensin 10 µM. **D.** Time course of the efflux of [^3^H]GABA in the presence of different betaine concentrations (1-100 mM). **E.** Time course of the efflux of [^3^H]GABA in the presence of monensin 10 µM and different betaine concentrations (1-100 mM). **F.** The kinetic analysis of the [^3^H]GABA-induced efflux with and without monensin 10 µM. **G.** Time course of the efflux of [^3^H]GABA in the presence of tiagabine 10 µM with and without monensin 10 µM. Data were fitted using Hill’s model and values are shown in the table at the bottom. Data are mean ± SEM from three individual experiments, performed in duplicate.

### Molecular dynamics and docking experiments show that betaine binds in the same binding pocket of GAT1 as GABA

In the absence of a crystal or cryo-EM structure of hGAT1 in the outward-facing conformation, we selected the Alphafold homology model to analyse the stability of bound GABA and betaine [34, 41, 42]. The bound ions (two Na^+^ and one Cl^-^) were added to the hGAT1 model using as a reference the outward-open human SERT crystal structure (PDB ID: 5I71) [43]. The docking studies for betaine to the outward-open hGAT1 provided a successful docking with the best fitness score of 42.24 (Figure 5 A). As a positive control, docking simulations for GABA were also run, which resulted in fitness score of 50.52 (Figure S5). The carboxyl head of betaine, which overlaps with the carboxyl head of docked GABA, interacts with Na1 (the sodium bound to the NA1 sodium binding site) and formed short-range (d < 3 Å) contacts with the backbone amide protons of L64 and G65, and with the side chain hydroxyl proton of Y140. The tail containing three methyl groups formed two medium-range (3 Å < d < 5 Å) contacts with residues S295 and S396.

**Figure 5:**
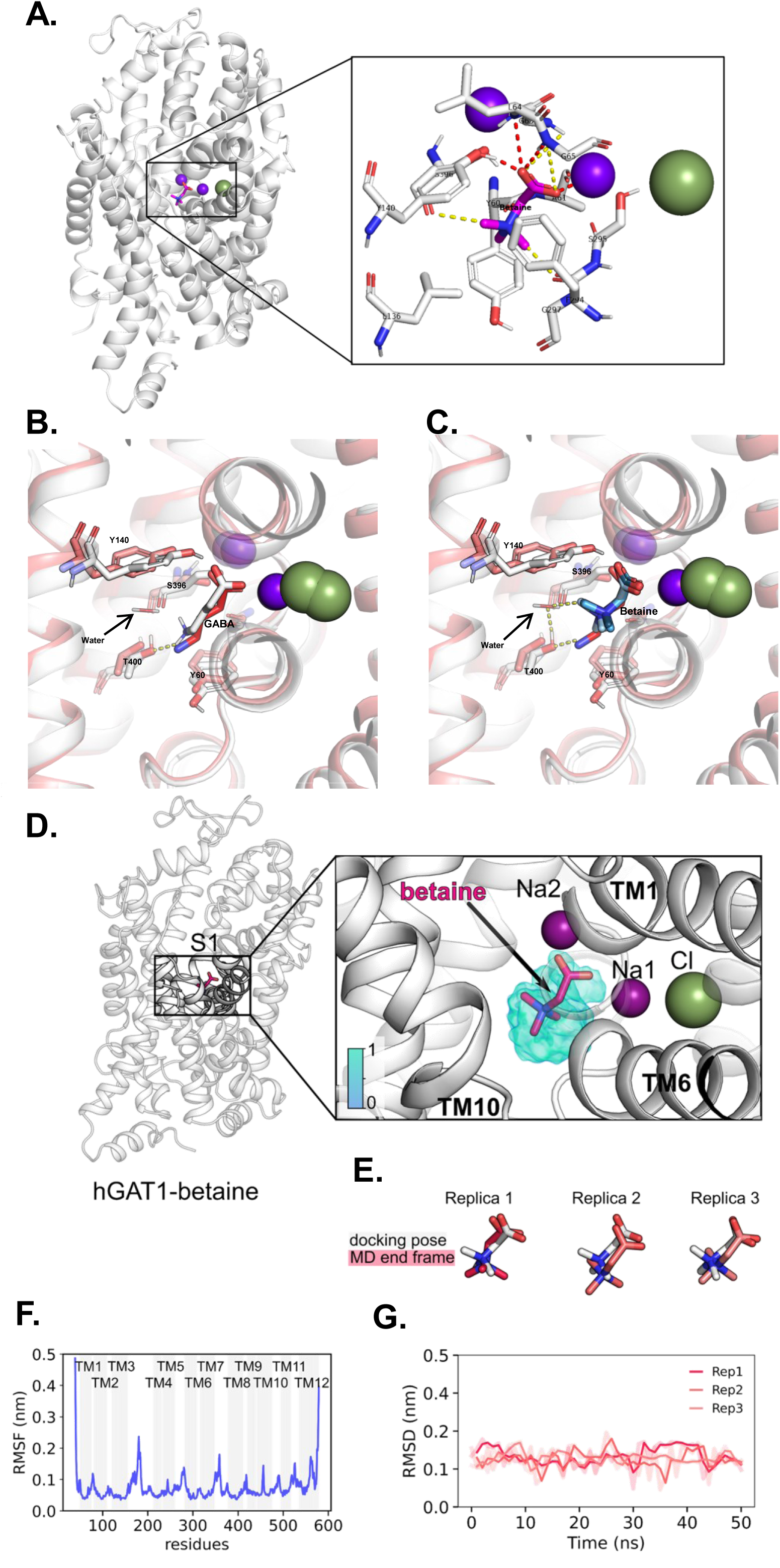
Molecular docking and MD simulation of betaine and GABA in hGAT1 show that betaine stably binds to GAT1 and forms less polar contacts than GABA. A. The successful docking of betaine in hGAT1 with zoomed-in view of the binding site. **B.** The overlapping of GABA-bound hGAT1 Alphafold in outward-open (in white) with cryo-Em structure of the hGAT1 in the inward-occluded (PDB: 7Y7W, in red) with zoomed-in view of the GABA binding site, in the presence of water molecule stabilized by T400 shown in yellow dashed line. **C.** The same overlapping for betaine-bound structure shows the tail of betaine forming hydrogen-bond with water molecule that is stabilized by carboxyl head of T400 shown in yellow dashed line. **D.** MD simulation results for hGAT1 in the outward open conformation bound to betaine is with a zoomed-in view as the S1-site. The average occupancy from three simulations is visualized using an isosurface, color-coded according to the legend. **E.** The docking poses (in white) and the respective end structures (in light red) at 50 ns resulting from MD simulations. **F.** The root mean square displacement of each replica smoothened with a running average over 2 ns. **G.** The root mean square fluctuation of hGAT1 residues by plotting a mean root mean square fluctuations value from the three replicas, emphasizing residues belonging to TM helices with a grey bar. In panel A, the short-range contacts (d < 3 Å) are indicated as red dashed lines and medium-range contacts (3 Å < d < 5 Å) as yellow dashed lines. The representation illustrated includes hGAT1 as ribbons (in panel A, B, C, D: outward-open in white. In panel C: inward-occluded in red), betaine and GABA shown as sticks, and Na^+^ (purple) and Cl^-^ (green) as spheres.

Interestingly, the recent work of Zhu et al. assigned five water molecules in GABA binding pocket in hGAT1. They showed in the inward occluded state, the amino acid group of GABA is stabilized by the hydroxyl group of T400 and a water molecule [44]. When we overlapped our hGAT1 model with GABA docked in the outward-open state with the inward-occluded cryo-EM structure of hGAT1 with GABA from Zhu et. al. (Figure 5 B), we observe similar stabilization of docked GABA in our model. Moreover, we also performed the same overlapping of the cryo-EM structure by Zhu et. al., with our outward-open hGAT1 with betaine docked (Figure 5 C). This overlapping showed that betaine molecule is not long enough to have a stable polar-contact with T400, but in the presence of a water molecule (stabilized by T400) it can form a stable hydrogen-bond and could allow the conformational change to the inward-open state.

The MD simulations were performed using the three docking poses of betaine with the best fitting scores (Figure 5 D, E). The simulations were run for 50 ns (the simulation parameters are reported in supplementary information as Code S1) and resulted in betaine stably bound to hGAT1, with root mean square displacement lower than 0.2 nm (Figure 5 F). The simulated GAT1 also resulted in a stable conformation, showing root mean square fluctuations greater than 0.2 nm only in residues belonging to the extracellular and intracellular loops (Figure 5 G).

### The relationship of GABA and betaine in rGAT1 depends heavily on their extracellular concentrations

As shown above (Figure 1, 2), betaine is a low affinity, secondary, and slow substrate of rGAT1. To further investigate the impact of betaine on GABAergic pathways, it is important to understand its relationship with GABA, the primary substrate of GAT1. We investigated the GABA-betaine relationship using TEVC and LC-MS/MS. The competitive assay experiments on TEVC were performed using six GABA concentrations ranging from 1-300 µM with twelve betaine concentrations from 0.001- 50 mM. The representative traces of the currents collected in the competitive assay at the holding potential of -60 mV were reported in Figure 6 A. With GABA concentration below 10 µM and betaine below 10 mM, we observed an inhibition of the transport current. With betaine 10 mM and above, the blocking effect disappeared, and a collective larger inward transport current was observed. The mean values of the currents for all 84 conditions were reported as a heat map of GABA-betaine competition (Data available in Table S3). The data showed that the GABA-betaine relationship starkly depends on their extracellular concentration (Figure 6 B).

**Figure 6:**
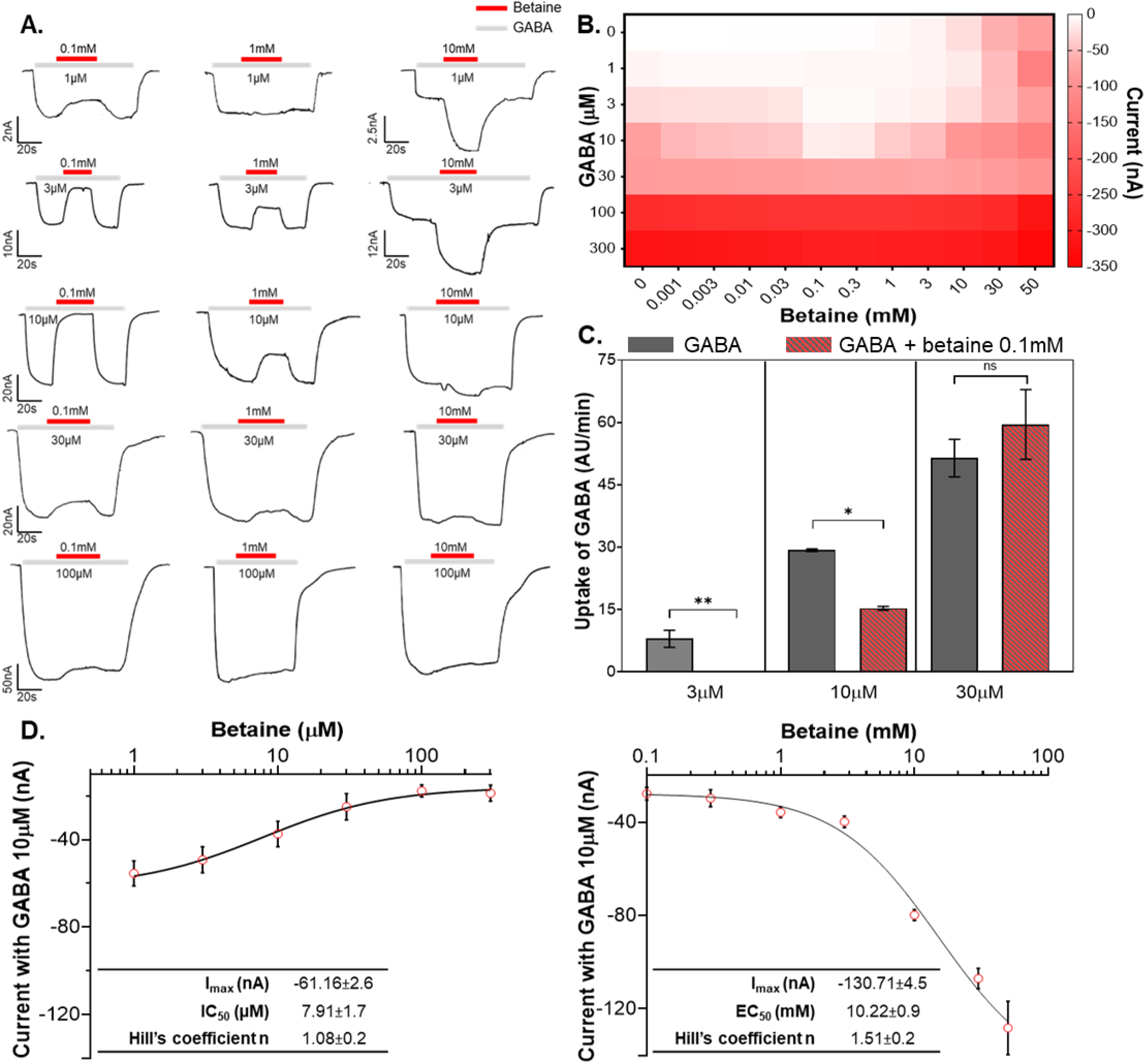
Betaine has a dual role in rGAT1, a GABA inhibitor at low concentrations and a secondary substrate at higher concentrations. A. Representative traces of GABA betaine assay in *X. laevis* oocytes expressing rGAT1 at holding potential V_h_ = -60 mV, where the oocyte was perfused with different GABA concentrations (1, 3, 10, 30, 100 μM) along with betaine 0.1, 1, and 10 mM. **B.** The heatmap analysis of the GABA betaine competitive assay shows their concentration-dependent relationship, using the combination of GABA (1-300 μM) with betaine (0.001-50 mM). Data are shown as mean ± SEM of 6/2 n/N. **C**. Detection of GABA and betaine in the oocytes incubated in GABA 3, 10, 30 µM with betaine 0.1 mM, using LC-MS/MS protocol. The qualitative analysis of GABA and betaine uptake by the oocytes is represented in this bar plot with the uptake values, as arbitrary units, of each oocyte per minute, data shown with SEM and obtained from n=3 with five oocytes in each sample. The p values were obtained by ordinary one-way ANOVA method followed by Bonferroni’s multiple comparisons test, with a single pooled variance with statistical significance of p<0.05. **D.** The kinetic analysis of different betaine concentrations (0.001-50 mM) with GABA 10 μM shows the dual behaviour of betaine in rGAT1, as at the lower concentrations (left) the GABA transport current is reduced with an increase in betaine, and at the higher concentration (right) the total transport current increases. All data were fitted using the logistic fitting model, fitting values shown in the inset, and current values shown as mean ± SEM of 6/2 n/N.

Since the detection time for GABA and betaine uptake by rGAT1 in LC-MS/MS protocol are different with distinct production-ions, it was possible to perform GABA and betaine competition in the same oocyte. Based on the electrophysiological findings, betaine shows the highest blocking of GABA transport 0.1 mM. Therefore, we incubated the oocytes expressing rGAT1 with GABA 3, 10, and 30 µM with betaine 0.1 mM for 25 minutes (n=3, five oocytes per sample). The cytosolic content detection by LC-MS/MS showed that in the presence of betaine 0.1 mM: with GABA 3 µM no GABA was taken up, with GABA 10 µM the GABA uptake was significantly reduced, and with GABA 30 µM the GABA uptake didn’t alter significantly, compared to the absence of betaine (Figure 6 C).

Taken together, these data showed that betaine has a dual effect on GABA transport by rGAT1. This duality can be best visualized at GABA 10 µM with different betaine concentrations (Figure 6 D). When GABA 10 µM is co-applied with betaine from 1-300 µM, concentration-dependent inhibition of GABA 10 µM transport current was observed. At the same GABA concentration, the perfusion with betaine from 1-50 mM yielded a concentration-dependent increase in transport current. This kind of dual effect of betaine on GABA transport vanished when the extracellular GABA concentration was 30, 100, and 300 µM i.e., larger than K_0.5,GABA_.

### Betaine slows down the rGAT1 transport cycle, denying binding of GABA that results in inhibition of GABA uptake by GAT1

The competition experiment of GABA and betaine was also performed with the voltage steps protocol to study steady and PSS transport currents. By observing the representative traces in the presence of GABA 10 µM with and without betaine 0.1 mM, the peculiar inhibitory action of betaine is visible. We observed a reduction in the current amplitude (indicated by the red dotted line) and increment in the decay time of the PSS currents in response to the voltage jumps (Figure 7 A). The I-V relationship of the current induced by GABA 10 µM in the absence and presence of betaine 0.1 mM revealed the blocking effect of betaine (Figure 7 B). Also, the Q-V analysis showed that in the presence of betaine 0.1 mM, the total charge dislocated by GAT1 did not increase significantly above that of GABA 10 µM alone (Figure 7 C). While the τ-V relationship for betaine 0.1 mM (Figure 2) showed an overall similar decay time constant as ND98, it became much faster for GABA 10 µM. However, τ for GABA 10 µM with betaine 0.1 mM, slowed down significantly, especially at the positive transmembrane potentials (Figure 7 D). Similarly, both decay constants, α and β for GABA 10 µM in the presence of betaine 0.1mM, showed an overall reduction, significantly in α at the positive transmembrane potentials (Figure 7 E). Altogether, it is evident that in the presence of betaine the transport rate of GAT1 for GABA decreased.

**Figure 7:**
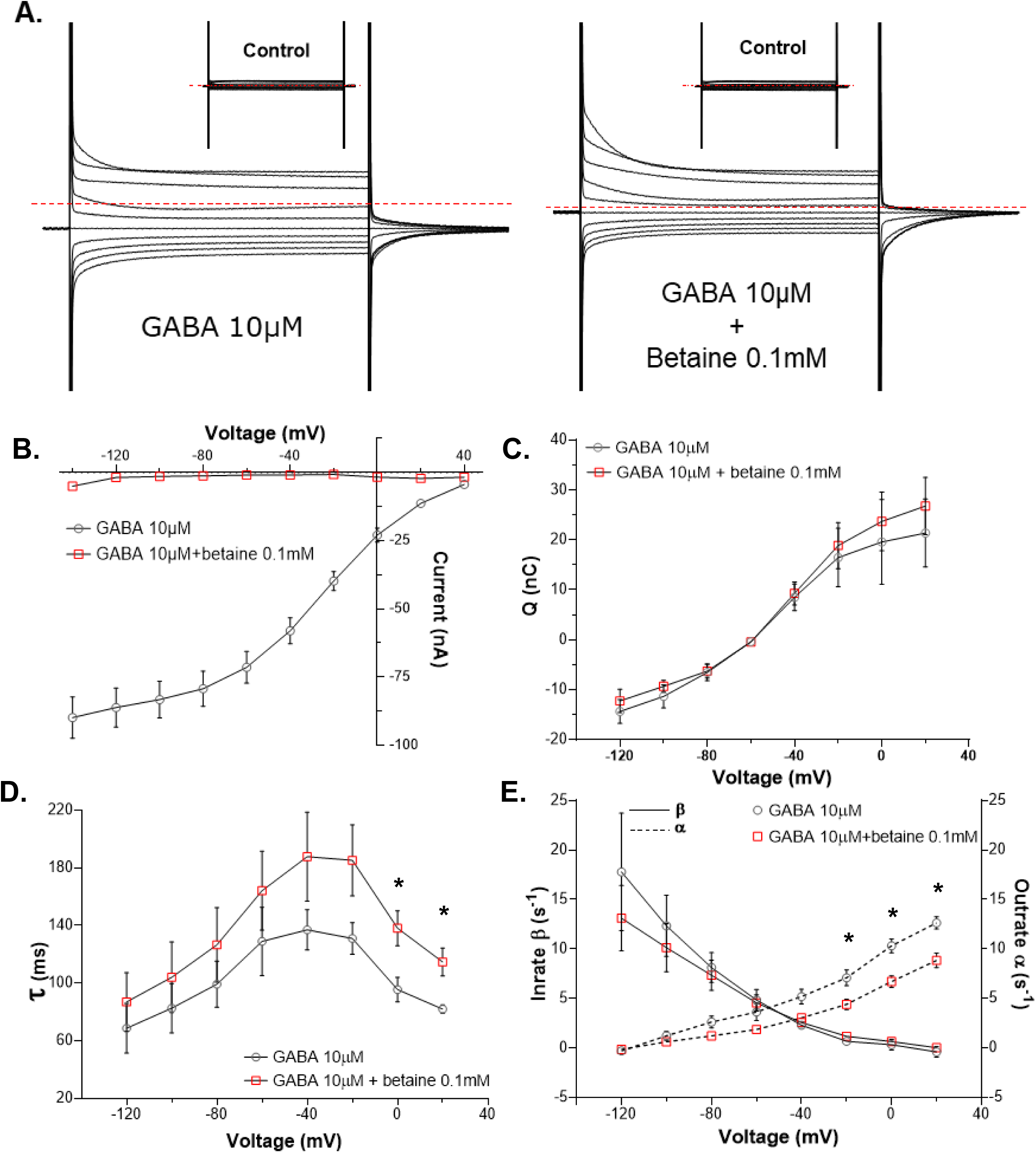
Betaine inhibits the GABA uptake by slowing down the transport cycle of rGAT1. **A.** The representative traces of the voltage-step response, from –140 mV to +40 mV, of the oocyte expressing rGAT1, at holding potential V_h_ = -60 mV, with non-saturating GABA 10 μM (left) and GABA 10 μM + betaine 0.1 mM (right), the dashed red line indicates the holding current for the oocyte at the holding potential. **B.** The current and voltage relationship of GABA 10 μM alone and with betaine 0.1 mM, from –120 mV to +20 mV, shows the transport current reduction at all voltages. **C.** The total charge dislocation and voltage relationship of GABA 10 μM alone and with betaine 0.1 mM, from –120 mV to +20 mV shows more charge dislocation happening in the presence of betaine. **D.** The total decay time constant and voltage relationship of GABA 10 μM alone and with betaine 0.1 mM, from –120 mV to +20 mV shows slowing down of the transport cycle in the presence of betaine. **E.** The relationships of unidirectional rate constants outrate (α) and inrate (β, shown as dashed line) with voltage for GABA 10 μM alone and with betaine 0.1 mM, from –120 mV to +20 mV. β in the presence of betaine does not decrease, but α decreases significantly at positive membrane potentials. All current responses were collected by giving 0.8 s long squared pulse at –20 mV of voltage jump. All current values shown as mean ± SEM of 3/1 n/N. The p values were obtained by the two-tailed p test with statistical significance of p<0.05.

## Discussion

Betaine is an endogenous molecule that plays an essential dual role as an osmolyte and a methyl donor in human physiology. In the past few decades, several studies have reported the beneficial effects of betaine in neurological and neurodegenerative diseases. Its systematic supplementation showed only mild side effects [4, 45, 46], making it a therapeutic molecule of interest. However, the presence of betaine in the brain and the mechanism involved leading to its positive action still remained a puzzle, deeming it a “dark knight” [4]. In this work, we show that betaine can be transported in a concentration-dependent manner by the most expressed neuronal GABA transporter GAT1, and that betaine modulates the uptake of GABA. From this data, we infer that betaine can modulate the extracellular GABA concentration and play a role in maintaining the E/I balance in the CNS.

In humans, mitochondria can produce betaine small quantities, but the primary source is the oral uptake from the dietary sources [1, 47]. As betaine is an excellent osmolyte, it tends to get accumulated in cells and tissues [48], leading to the underestimation of betaine absorption when using the traditional blood plasma assays [49, 50]. The pharmacokinetics of this accumulation is dose-dependent with the peak betaine concentration reaching 1-3 mM within hours after intake [51, 52]. In liver and kidney, the concentration of betaine can reach up to ∼30 mM and ∼100 mM respectively [4, 53]. The reported betaine level in the mice brain is unclear [54], although recently Knight et al showed that the neurons from the hippocampal tissues of mice, can uptake and accumulate betaine up to 12 mM [50]. From their findings, it is conceivable that the neuronal concentrations of betaine might reach mM levels. With this argument along with the fact that the affinity of betaine for BGT-1 also ranges towards millimolar [26], we used betaine at millimolar levels to study its interaction with GAT1 and BGT- 1.

The electrophysiological measurements using TEVC in *X. laevis* oocytes heterologously expressing rGAT1 resulted in betaine-induced concentration-, voltage-, and sodium-dependent inward transport currents, which could be inhibited by GAT1 inhibitors like SKF89976a (Figure 1), tiagabine, and NO-711 (Figure S1). The transport of betaine by GAT1 was also confirmed with automated patch-clamp (using the Patchliner™) on CHO cells overexpressing hGAT1 at V_h_ = -80 mV with EC_50_ ≈ 27 mM (Figure S1). We also showed betaine translocation by GAT1 using radiolabelled efflux assay and LC-MS/MS analysis of the cytosol contents of *X. laevis* oocytes (Figure 3 and 4).

Interestingly, the kinetic analysis of the GABA and betaine transport currents in GAT1 and BGT-1 reveal something interesting (Table in Figure 1). For GABA, the transporter efficiency (I_max_/K_0.5_) of GAT1 is about ten times larger than BGT-1, meaning that GAT1 can transport a lot more GABA than BGT-1. Whereas, for betaine, the transporter efficiency of GAT1 is half of BGT-1. This suggests that compared to GABA there is little difference in the amount of betaine that can be transported by GAT1 and BGT-1. Thus, BGT-1 cannot transport GABA efficiently like GAT1, but GAT1 can transport betaine with efficiency close to that of BGT-1.

The detailed analysis of the PSS currents gives information about the initial steps of the transporter cycle and the interaction of the transporter with ions and substrate [22]. The analysis of the decay time constant (τ) of the large transient currents as inrate β and outrate α constants showed that in the presence of betaine (0.1 and 1mM), β of rGAT1 increased. This implies that when betaine is present at low concentration and binds in the GAT1 vestibule, it cannot induce the same fast conformational changes, as GABA [37]. Also, the higher inrate than the outrate of charge dislocation suggests that in the presence of betaine, especially at lower concentrations, the transporter cannot transit to the inward-open state as fast as in the presence of GABA. This behaviour of GAT1 suggests that it requires a third “interaction” to transport betaine as in BGT-1 [26] or that the conformational transition towards the inward open state is less efficient. This is further supported by the Q-V relationship of betaine in GAT1, where the curve is shifted towards more positive voltage (for 0.1 mM betaine: V_0.5_ = - 27.16 ± 2.57 mV, voltage where half of the charge is moved) than for ND98 alone (V_0.5_ = -32.87 ± 1.10 mV) and GABA (for GABA 10 µM V_0.5_ = -43.19 ± 3.38 mV) (Figure 7 C and Table S1). A similar shift in V_0.5_, could also occur when the extracellular Na^+^ concentration is changed [30]. In the experiments reported here, extracellular Na^+^ concentrations were unaltered, consequently, this shift can be related to either an apparent change in Na^+^ concentration inside the transporter vestibule or a decrease of the Na^+^ dissociation [55]. Moreover, the molecular docking and MD simulations of betaine-bound hGAT1 showed that it forms less short-range contacts than GABA (Figure 5 A, S5). This implies that it would require involvement of more residues to stabilize the binding of betaine than GABA, suggesting it as a factor behind the lower affinity for betaine in GAT1. In summary, the transition of GAT1 out of the occluded state is slower for betaine than GABA, but it might not be the only possible interpretation of our data.

The GABA-betaine competitive assay experiments clearly showed that their relationship is strongly concentration-dependent (Figure 6). At high GABA concentrations, betaine shows minimal impact on the GABA transport by GAT1. In contrast, when the extracellular GABA concentration is lower than its K_0.5_ (≈16 µM), betaine at concentrations of 0.1-3 mM inhibits GABA transport by GAT1. The reduction of the transport rate in the presence of betaine has a role in this selective inhibition of GABA transport. In the presence of low extracellular betaine, both the inrate and outrate of GABA reduced starkly (Figure 7 D, E and Table S2). As the time required for betaine to be moved inside the cell is longer than GABA, the betaine remains bound to some of the GAT1 proteins, reducing the available transporters able to complete the cycle in the presence of GABA lower than K_0.5,GABA_. Consequently, several transporters fail in rapid completion of their cycle, enabling the modulation of the GABA uptake by betaine. When the concentration of GABA reaches close to the saturation, it outcompetes betaine allowing rapid completion of the transport cycle. While at high concentrations (>K_0.5,betaine_), betaine behaves like a regular substrate as the higher chemical gradient provides energy for its slower intracellular translocation by GAT1. This inhibitory role of betaine was also confirmed by LC-MS/MS, which showed that the uptake of extracellular GABA, when <K_0.5,GABA_, was significantly reduced by betaine 0.1 mM (Figure 6 C). This kind of selective inhibition by betaine appears to be specific to GAT1 and was not observed in BGT-1 (Figure S6). Also interestingly, betaine did not show any selective inhibition of the nipecotic acid (another GAT1 substrate) induced transport currents in GAT1 (Figure S7).

The conventional GAT1 blockers derived from its competitive inhibitor nipecotic acid were modified with lipophilic sidechains to allow the blood-brain barrier (BBB) penetration and disable the substrate properties [56, 57]. One such derivative, tiagabine, which is the only anticonvulsant drug targeting GAT1, is used to treat partial seizures in patients with epilepsy. However, tiagabine treatment has been reported to cause several unwanted side effects such as confusion, abnormal mood swings, dizziness, tremors, fatigue, and nervousness [58]. The recent works on GAT1 structures suggest that tiagabine may bind GAT1 in the outward open conformation and blocks in the inward open conformation, disabling any further uptake by the transporter [35, 36, 44, 59]. Differently from tiagabine, betaine slows down the transport cycle of GAT1 and triggers a temporal inhibition of GABA uptake. This effect can be abrogated by the increase of extracellular GABA, which will allow the transporter to regain its original function of the GABA uptake. Along with this unique blocking characteristic and the minimal harmful side-effects associated with its supplementation, betaine could be a useful molecule to modulate GABA homeostasis targeting GAT1.

Importantly, betaine does not require chemical modifications to cross the BBB. Wang et al. showed *in vivo* that the cylindrical polymer brushes (CPB) modified with betaine could cross the BBB [60]. They also performed *in vitro* uptake experiments on endothelial bEnd.3 cells to identify the transport mechanism for the modified CPBs and deemed BGT-1 as the transporter responsible for the translocation. Interestingly, the uptake inhibition of the modified CPB by GABA was much stronger than by betaine, which they correlated with the higher affinity of BGT-1 for GABA than betaine. By taking in account our results of betaine transport by GAT1, it would suggest that along with BGT-1, also GAT1, which is expressed in bEnd.3 cells [61], could have contributed to the translocation of the modified CPBs.

The E/I balance in the CNS is essential for the healthy, stable, and physiologically functional neuronal circuits; and the disruption of this balance is the primary cause of many neurological diseases [62]. Given that GABA is the primary inhibitory neurotransmitter in the CNS, its homeostasis is essential for the formation of learning and memory. The systematic supplementation of betaine has been shown to ameliorate the E/I balance-related diseases such as Alzheimer’s, Parkinson’s, dementia, schizophrenia, and stress-related disorders, but these results lack substantial mechanistic explanation [4]. Since the extracellular GABA levels are critical for these disorders and diseases, the peculiar GABA inhibitory behaviour of betaine can explain its beneficiary effects from the molecular point of view.

Recently Knight et al. showed that the hippocampal neurons not only uptake and accumulate betaine, but also modulate the neuronal uptake of other essential osmolytes in response to the changes in osmolarity [50]. Especially under hyperosmotic conditions, hippocampal neurons preferentially uptake betaine over glycine and glutamine. This connects neuronal uptake of betaine with glycine, an essential inhibitory neurotransmitter, and glutamine that is the precursor to the production of both glutamate (the primary excitatory neurotransmitter) an GABA (the primary inhibitory neurotransmitter). Such interaction of betaine with different essential elements of neurophysiological processes suggests its participation in more than one [63]. Especially, with the GABA inhibitory behaviour of betaine shown here suggests a possibly larger modulatory role of betaine in maintaining the E/I balance in the CNS. Moreover, it was shown by Hardege et al. that the *Caenorhabditis elegans* nervous system can synthesize betaine, specifically in the interneurons, and use it as a modulator of different behavioural states [21]. They also showed that the vesicular transporter CAT-1 expressed in the interneurons could transport betaine, suggesting nematode neurons can load betaine in the synaptic vesicles. Lastly, the work from Kunisawa et al. showed that the lack of betaine in rats could affect GABAergic transmission and memory formation [19]. They also demonstrated that these effects of betaine are regulated not solely by BGT-1 but partly mediated by betaine modulating the GABAergic system. These results along with the data presented here, strongly suggest that betaine should have a specific role in the CNS (possibly a neuromodulator) apart from being an osmoregulator and a methyl donor.

Overall, this work confirms the betaine transport by GAT1 and the concentration-dependent modulation of GABA transport by GAT1 using betaine. Naturally, these findings raise the question of whether other neurotransmitter transporters and receptors may interact with betaine. While our work answers one of the questions around the role of betaine, it is now even more important to explore its presence and investigate its effects in the brain, beyond being a substrate of BGT-1. Betaine carries the potential of being a natural neurotherapeutic agent and given its already an FDA-approved drug (Cystadane^®^ for homocystinuria), it would be of great interest to explore its applications in treatments targeting the GABAergic system in general.

## Online Methods

### Experimental Model and Subject Details

#### Cell lines

Full-grown *Xenopus laevis* oocytes were maintained at 18°C in post-injection NDE solution (96 mM NaCl, 2 mM KCl, 1 mM MgCl_2_, 5 mM HEPES, 2.5 mM pyruvate, 0.05 mg/mL gentamicin sulphate, and 1.8 mM CaCl_2_ at pH 7.6).

Human embryonic kidney 293 (HEK293) cells stably expressing the rat isoform of GAT1 (rGAT1) were used (rGAT1-HEK293 cells) in batch release assays. The generation and maintenance of stable cell lines expressing rGAT1 was conducted as previously described by other for other SLC6 transporters [64]. rGAT1-HEK293 cells were maintained in high glucose (4.5 g/L) and L-glutamine-containing DMEM, supplemented with 10% FBS and G418 (250 µg/mL) in a humidified atmosphere (37°C, 5% CO_2_) and in a sub-confluent state.

CHO cells were grown and maintained in DMEM containing L-glutamine (PAA laboratories, Austria) with gentamicin 50 mg/L, geneticin 500 µg/mL, 10% foetal calf serum on 100 mm-diameter cell culture dishes at 37°C in an atmosphere of 5% CO_2_ and 95% air [65].

### Method Details

#### Heterologous Expression in *X. laevis* oocytes

The detailed experimental procedure has been described elsewhere [55]. Briefly, the cDNAs encoding rat GAT1 in the pAMV vector, and Madin-Darby canine kidney (MDCK) canine BGT-1 in the pSPORT1 vector were linearized using NotI, and the in vitro cRNA synthesis was done in the presence of Cap Analog 10 mM and T7 RNA polymerase 200 U.

Oocytes were collected from *X. laevis* female frogs, under anaesthesia, by performing laparotomy on the abdomen, to remove portions of the ovary. Detailed procedures are described elsewhere [25]. For the heterologous expression of the GABA transporters, stage IV oocytes were micro-injected with cRNA encoding the target protein i.e., rGAT1 and cBGT-1 (12.5 ng/ 50 nL) and incubated for the expression period (3-4 days) at 18°C before the experimental procedure.

The oocytes used in this work were either bought from EcoCyte Bioscience GmbH, Germany or donated by a collaborator with approved experimental protocol by the Committee of the “Organismo Preposto al Benessere Animale” of the University of Insubria, Varese, Italy, and nationally by Ministero della Salute (n. 449/2021-PR), Italy; and used in agreement with the Art.18 of decreto legislativo 4 marzo 2014, n. 26 and Art. 26 in directive 2010/63/eu of the European parliament and of the council of 22 September 2010.

#### Electrophysiology

The two-electrode voltage clamp (TEVC) Oocyte ClampOC-725B, Warner Instruments, Hamden, CT, USA was used for controlling voltages and recording currents. The clamped oocytes were maintained under continuous perfusion of the basic buffer solution ND98 (NaCl 98 mM, CaCl_2_ 1.8 mM, MgCl_2_ 1 mM, HEPES 5 mM at pH7.6) with a flow rate of 1.5 mL/min. For dose-response curves and competitive assay experiments, solution exchange was performed using an in-house developed gravity-driven digitally controlled perfusion control system. Signals were filtered at 0.1 kHz and sampled at 1 kHz. The voltage-jump experiments were performed by the voltage pulse control protocol consisting of 10 square pulses (0.8 s long) from -140 mV to +40 mV (at 20 mV increments). All recorded currents were digitalized using Axon CNS 1440B Digidata system controlled by pClamp 11.2.2 (Molecular Devices, USA).

For whole-cell patch clamp experiments using CHO cells expressing human GAT1 on the Patchliner™ (Nanion Technologies GmbH, Germany), the cell-harvesting was performed right after the setup of the instrument, to be transferred directly into the CellHotel^TM^. The cells were automatically transferred from the CellHotel^TM^ to each of the eight wells of 4X chips for Patchliner™ that was set in the whole-cell mode with the holding membrane potential at -80 mV.

The internal solution (10 mM NaCl, 110 mM KF, 10 mM EGTA, 10 mM KCl, and 10 mM HEPES at pH 7.2) was provided inside the cells, followed by the external solution outside (Ca^+2^-rich, 130 mM NaCl, 4 mM KCl, 10 mM CaCl_2_, 1 mM MgCl_2_, 5 mM D-Glucose monohydrate, 10 mM HEPES at pH 7.4) for seal enhancement. For the recording on the Patchliner™, only cells showing a seal resistance >300 MΩ were taken into consideration for the analysis. After the seal was established, a double wash with external solution (standard, 140 mM NaCl, 4 mM KCl, 2 mM CaCl_2_, 1 mM MgCl_2_, 5 mM D-Glucose monohydrate, 10 mM HEPES at pH 7.4) was performed to remove the extra Ca^2+^. The following fluidic protocol was followed for all betaine concentrations: 6 s perfusion of 60 µL of betaine-containing external solution (speed 10 µL/s) followed by 18 s perfusion of 180 µL of external solution (standard) to wash away the betaine (speed 10 µL/s).

#### Substrate detection in *X. laevis* oocytes using LC-MS/MS

To perform the substrate uptake experiment, the *X. laevis* oocytes expressing rGAT1 were incubated in a fresh Petri dish containing the substrate+ND98 for 25-30 minutes at 18°C. To remove any presence of the extracellular substrate post-incubation, the oocytes were carefully washed thrice with ice-cold ND96 for two minutes. At the end of the washing, the oocytes (in the group of 5) were transferred in a fresh Eppendorf containing CH_3_OH:CHCl_3_:H_2_O in proportion of 3:1:1 v/v with the solvent to oocytes ratio of 1:100. The oocytes were homogenized using water ultra-sonicator and then centrifuged at 13,000xg for eight minutes at 4°C. The supernatant from these samples was removed, and the pallet containing the cytosol content with oocyte debris were diluted in H_2_O (MilliQ, Merck, Italy) to extract the hydrophilic compounds. These samples were centrifuged at 13,000xg for eight minutes at 4°C and the aqueous top part containing cytosol content was extracted and used immediately for LC-MS/MS detection or stored at -80°C to use later (up to three months).

Betaine and GABA standard stock solutions (1 mM in 0.1% formic acid) were prepared in MS grade water. For quantitative mass spectrometry, a Finnigan LXQ linear ion trap mass spectrometer, equipped with an ESI ion source (Thermo Electron Corporation, CA, USA) was used. The analyses were performed in positive (spray voltage 4.5 kV, capillary temperature 270°C) and in the multiple-reaction monitoring (MRM) mode. The tuning parameters, the optimization of collision energy for each substance, and the choice of target compound fragments were conducted in continuous flow mode by using standard solutions at a concentration of 5 μM. The MRM acquisitions for GABA and betaine were accomplished by monitoring the 114/89 and 118/59 transitions, respectively. The HPLC analysis was performed using a Finnigan Surveyor MS plus HPLC system (Thermo Electron Corporation, CA, USA). For both GABA and betaine quantitation, separation was achieved using the C18 column (ACQUITY UPLC Peptide BEH C18 Column, 300 Å, 1.7 µm, 2.1 mm×150 mm). The mobile phase was composed of (A) water with 0.1% (v/v) formic acid and (B) acetonitrile (100%) plus 0.1% (v/v) formic acid with a flow rate 200 µL/min; gradient 0–7.0 min/2% (v/v) B, 7– 10 min/2–50% (v/v) B; 10-15 min/50-2% (v/v) B.

To verify the sensitivity and linearity of the LC-MS/MS response to GABA and betaine, calibration curves showing the relative peak areas plotted against analytes concentrations (range 2–16 µM) were created. GABA and betaine contents were expressed as relative intensities (peak areas) among samples.

#### Radiolabelled GABA release assay

The batch-release assays have been conducted as described before [66, 67]. In brief, the transfected HEK293 cells overexpressing rGAT1 were grown overnight onto poly-D-lysine coated 96-well plates (4×10^4^ cells/well). After removing DMEM, the cells were loaded with 0.01µM [^3^H]GABA in Krebs-Ringer HEPES buffer (KHB; NaCl 120 mM, KCl 3 mM, CaCl_2_ 2 mM, MgCl_2_ 2 mM, glucose 20 mM, HEPES 10 mM at pH 7.3) for 20 min (5% CO_2_, 37°C). After the incubation, the cells were brought to room temperature and washed five times with KHB during 15min. Afterwards, cells were kept during 10 min either in KHB or KHB + monensin (mon) 10 μM and the resulting supernatant was discarded. The batch-release assay started by applying either KHB or KHB + mon 10 μM (100 μL; every 2min, four times). Subsequently, the substance of the interest (in KHB or KHB + mon 10μM) was added at a specific concentration (every 2 min, five times). The resulting supernatants were collected and transferred to neighbour wells. Tiagabine (10 μM) and GABA (1-300 μM) were used as negative and positive controls, respectively. Three independent experiments were performed in duplicate for every compound and respective concentration of interest. At the end, liquid scintillation cocktail was added to the wells with remaining cells, with transferred supernatant and to the wells used for total uptake and activity measurements. The total radioactivity present in the supernatant and in the remaining cells was set as 100%, and the amount of [^3^H]GABA present in the supernatant was expressed as percentage of the total.

#### Molecular docking and molecular dynamics simulations

In the absence of a crystal or cryo-EM structure of GAT1 in the outward-facing conformation, we selected the Alphafold homology model to analyse the stability of the docked and bound betaine [41, 42]. The docking experiments were performed using GOLDscore scoring function on GOLD molecular docking software (The Cambridge Crystallographic Data Centre, UK). For MD simulations, the bound-ions were added to the GAT1 model using the outward-open human SERT crystal structure (PDB ID: 5I71) [43] as a reference. The generated system was converted from full-atom into a coarse grain representation using the MARTINI force field [68–70], with a membrane composition of 1-palmitoyl-2-oleoyl phosphatidylcholine (POPC): cholesterol containing membrane (POPC:CHOL 70:30 mol%) [71] solvated in water with 150mM NaCl. After 1μs of coarse-grain simulation, while restraining the protein structure, the membrane was equilibrated. The coarse-grained system was then converted to an all-atom representation [72] in which the transporter was replaced by the original GAT1 model to avoid factitious structural imprecisions induced by the double coordinate conversion of the protocol. Betaine was positioned according to the docking results. Possible atom overlaps between the reinserted protein and the relaxed membrane were loosened using the membed procedure [73] as previously described [74]. We used the amber ff99SB-ILDN force field[75] to describe GAT1, ions and the solvent, and Slipid [76] for POPC and cholesterol. Previously reported structural analysis of the orthologous SERT structure [43], suggests that the residue E467 is protonated. All simulations were carried out with GROMACS version 2021.4 [77]. Three replicas of the final assembled system were energy-minimised and equilibrated in four steps of 2.5 ns, each by stepwise releasing the position restraints (1,000, 100, 10, 1 kJ/mol/nm) that are active on the Cα atoms and the bound chloride ion [78, 79]. Per each trajectory, the production run was carried for 500ns after removing all position restraints. The temperature was maintained at 310 K using the v-rescale (τ = 0.5 ps) thermostat [80], while separately coupling protein, membrane, and solvent. The pressure was maintained at 1 bar using the Parrinello-Rahman barostat [81] in a semi-isotropic manner and applying a coupling constant of 20.1 ps. Long-range electrostatic interactions were described using the smooth particle mesh Ewald method [82] applying a cutoff of 0.9 nm. The van der Waals interactions were described using the Lennard Jones potentials applying a cutoff of 0.9 nm. Long-range corrections for energy and pressure were applied. Coordinates of all atoms were recorded every 5 ps. The complete set of parameters of the production run can be found in the supplementary materials (Code S1).

### Quantification and statistical analysis

#### Electrophysiological analysis

For TEVC, data analysis was performed using Clampfit 11.2 (Molecular Devices, USA). The dose-response currents were calculated as the maximum current amplitude during the perfusion, and the curves were fitted by non-linear regression to the Logistic equation using Originpro 8 (OriginLab Corporation, USA)..The pre-steady state currents in rGAT1 were isolated by subtraction of the corresponding trace in the presence of SKF89976a 30µM, as this molecule blocks all of the transport-associated steady state and transient currents [37, 83]. The resultant traces were fitted with single exponentials to obtain decay time constant and voltage (τ-V) curves. The same traces were used to obtain the charge dislocation-voltage (Q-V) curves by fitting with Boltzmann equation to the sigmoidal curve, which also provided maximal displaceable charge Q_max_ [37, 38]. The further derivation of τ-V and Q-V curves yielded the unidirectional rate constants inrate (α) and outrate (β) (Eq. 1).

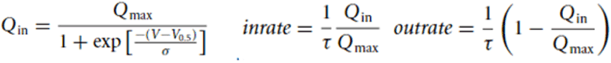

Equation 1: The Boltzmann equation quantifies Q_in_, total displaced charge. Where Q_max_ is the maximal displaceable charge, V_0.5_ is the voltage at which half maximal charge is displaced (i.e., midpoint of the sigmoidal), σ represents the slope factor. The inrate and outrate constants provide the unidirectional constants α and β respectively.

The data collected with Patchliner™ were saved as ASCII files and successively analysed with the software OriginPro 2021 (OriginLab Corporation, USA). The traces were filtered (using a weighted Adjacent-Averaging filter with ten points of window) and for each cell the baseline was subtracted to the respective betaine traces. The transport current was read as average ± SD (standard deviation) of the current value between 4-6 seconds after betaine perfusion. It is to be noticed that the current in each well is generated by the sum of currents coming from four cells simultaneously since each well of a 4X chip contains four holes for cell attachment.

## Acknowledgements

We thank Dr. Raffella Cinquetti for the technical support during membrane protein expression in *X. laevis* oocytes.

## Statements & Declarations

### Author Contributions

**Manan Bhatt:** Formal analysis and investigation, conceptualization, methodology, validation, visualization, writing – original draft preparation, writing – review and editing. **Erika Lazzarin:** Formal analysis and investigation, validation, visualization, writing – review and editing. **Ana Sofia Alberto-Silva:** Formal analysis and investigation, validation, writing – review and editing. **Guido Domingo:** Formal analysis and investigation, validation, methodology, writing – review and editing. **Rocco Zerlotti:** Formal analysis and investigation, visualization, writing – review and editing. **Ralph Gradisch:** Formal analysis and investigation, validation. **Andre Bazzone:** Supervision, funding acquisition, writing – review and editing. **Harald H. Sitte:** Supervision, funding acquisition, writing – review and editing. **Thomas Stockner:** Supervision, funding acquisition, conceptualization, writing – review and editing. **Elena Bossi:** Conceptualization, supervision, methodology, validation, funding acquisition, writing – review and editing.

### Consent for publication

Not applicable

### Materials availability

All unique/stable reagents generated in this study are available from the lead contact without restriction.

### Data Availability

The data that support the findings of this study are available from the corresponding author upon reasonable request.

### Ethics approval

This animal study was reviewed and approved by the Committee of the “Organismo Preposto al Benessere degli Animali” of the University of Insubria and nationally by Ministero della Salute (permit nr. 449/2021-PR).

## References

1. Lever, M. and S. Slow, The clinical significance of betaine, an osmolyte with a key role in methyl group metabolism. Clin Biochem, 2010. 43(9): p. 732–44.DOI: 10.1016/j.clinbiochem.2010.03.009.

2. Craig, S.A.S., Betaine in human nutrition. The American Journal of Clinical Nutrition, 2004(80): p. 539–549.DOI: 10.1093/ajcn/80.3.539.

3. Arumugam, M.K., et al., Beneficial Effects of Betaine: A Comprehensive Review. Biology (Basel), 2021. 10(6).DOI: 10.3390/biology10060456.

4. Bhatt, M., et al., Betaine - the Dark Knight of the brain. Basic Clin Pharmacol Toxicol, 2023.DOI: 10.1111/bcpt.13839.

5. Chai, G.S., et al., Betaine attenuates Alzheimer-like pathological changes and memory deficits induced by homocysteine. J Neurochem, 2013. 124(3): p. 388–96.DOI: 10.1111/jnc.12094.

6. Sun, J., et al., Association between malnutrition and hyperhomocysteine in Alzheimer’s disease patients and diet intervention of betaine. J Clin Lab Anal, 2017. 31(5).DOI: 10.1002/jcla.22090.

7. Ibi, D., et al., Involvement of GAT2/BGT-1 in the preventive effects of betaine on cognitive impairment and brain oxidative stress in amyloid beta peptide-injected mice. Eur J Pharmacol, 2019. 842: p. 57–63.DOI: 10.1016/j.ejphar.2018.10.037.

8. Alirezaei, M., et al., Beneficial antioxidant properties of betaine against oxidative stress mediated by levodopa/benserazide in the brain of rats. J Physiol Sci, 2015. 65(3): p. 243–52.DOI: 10.1007/s12576-015-0360-0.

9. Yoshihara, S., et al., Betaine ameliorates schizophrenic traits by functionally compensating for KIF3-based CRMP2 transport. Cell Rep, 2021. 35(2): p. 108971.DOI: 10.1016/j.celrep.2021.108971.

10. Qu, Y., et al., Betaine supplementation is associated with the resilience in mice after chronic social defeat stress: a role of brain-gut-microbiota axis. J Affect Disord, 2020. 272: p. 66–76.DOI: 10.1016/j.jad.2020.03.095.

11. Nie, C., et al., Betaine reverses the memory impairments in a chronic cerebral hypoperfusion rat model. Neurosci Lett, 2016. 615: p. 9–14.DOI: 10.1016/j.neulet.2015.11.019.

12. Li, Q., et al., Betaine protects rats against ischemia/reperfusion injury-induced brain damage. J Neurophysiol, 2022. 127(2): p. 444–451.DOI: 10.1152/jn.00400.2021.

13. Ohnishi, T., et al., Investigation of betaine as a novel psychotherapeutic for schizophrenia. EBioMedicine, 2019. 45: p. 432–446.DOI: 10.1016/j.ebiom.2019.05.062.

14. Cesar-Razquin, A., et al., A Call for Systematic Research on Solute Carriers. Cell, 2015. 162(3): p. 478–87.DOI: 10.1016/j.cell.2015.07.022.

15. Zhou, Y., et al., The betaine-GABA transporter (BGT1, slc6a12) is predominantly expressed in the liver and at lower levels in the kidneys and at the brain surface. Am J Physiol Renal Physiol, 2012. 302(3): p. F316–28. DOI: 10.1152/ajprenal.00464.2011.

16. Uchida, Y., et al., Quantitative targeted absolute proteomics of human blood-brain barrier transporters and receptors. J Neurochem, 2011. 117(2): p. 333–45.DOI: 10.1111/j.1471-4159.2011.07208.x.

17. Kempson, S.A., Y. Zhou, and N.C. Danbolt, The betaine/GABA transporter and betaine: roles in brain, kidney, and liver. Front Physiol, 2014. 5: p. 159.DOI: 10.3389/fphys.2014.00159.

18. Nishimura, T., et al., System A amino acid transporter SNAT2 shows subtype-specific affinity for betaine and hyperosmotic inducibility in placental trophoblasts. Biochim Biophys Acta, 2014. 1838(5): p. 1306–12.DOI: 10.1016/j.bbamem.2014.01.004.

19. Kunisawa, K., et al., Betaine attenuates memory impairment after water-immersion restraint stress and is regulated by the GABAergic neuronal system in the hippocampus. Eur J Pharmacol, 2017. 796: p. 122–130.DOI: 10.1016/j.ejphar.2016.12.007.

20. Rufener, L., et al., acr-23 Encodes a monepantel-sensitive channel in Caenorhabditis elegans. PLoS Pathog, 2013. 9(8): p. e1003524.DOI: 10.1371/journal.ppat.1003524.

21. Hardege, I., et al., Neuronally produced betaine acts via a ligand-gated ion channel to control behavioral states. Proc Natl Acad Sci U S A, 2022. 119(48): p. e2201783119.DOI: 10.1073/pnas.2201783119.

22. Bhatt, M., et al., A comparative review on the well-studied GAT1 and the understudied BGT-1 in the brain. Front Physiol, 2023. 14: p. 1145973.DOI: 10.3389/fphys.2023.1145973.

23. Zafar, S. and I. Jabeen, Structure, Function, and Modulation of gamma-Aminobutyric Acid Transporter 1 (GAT1) in Neurological Disorders: A Pharmacoinformatic Prospective. Front Chem, 2018. 6: p. 397.DOI: 10.3389/fchem.2018.00397.

24. Bi, D., et al., GABAergic dysfunction in excitatory and inhibitory (E/I) imbalance drives the pathogenesis of Alzheimer’s disease. Alzheimers Dement, 2020. 16(9): p. 1312–1329.DOI: 10.1002/alz.12088.

25. Bhatt, M., et al., The “www” of Xenopus laevis Oocytes: The Why, When, What of Xenopus laevis Oocytes in Membrane Transporters Research. Membranes (Basel), 2022. 12(10).DOI: 10.3390/membranes12100927.

26. Matskevitch, I., et al., Functional characterization of the Betaine/gamma-aminobutyric acid transporter BGT-1 expressed in Xenopus oocytes. J Biol Chem, 1999. 274(24): p. 16709–16.DOI: 10.1074/jbc.274.24.16709.

27. Forlani, G., et al., Three kinds of currents in the canine betaine-GABA transporter BGT-1 expressed in Xenopus laevis oocytes. Biochim Biophys Acta., 2001. 1538(2-3): p. 172–80.DOI: 10.1016/s0167-4889(00)00144-0.

28. Guastella, J., et al., Cloning and expression of a rat brain GABA transporter. Science, 1990. Sep 14;249(4974). DOI: 10.1126/science.1975955.

29. Schousboe, A. and P. Krogsgaard-Larsen, SKF89976A, A Highly Potent GABA Transport Inhibitor Capable of Crossing the Blood–Brain Barrier, in Reference Module in Biomedical Sciences. 2018, Science Direct: Biomedical Sciences DOI: 10.1016/B978-0-12-801238-3.97480-4.

30. Mager, S., et al., Ion binding and permeation at the GABA transporter GAT1. J neurosci, 1996. 16(17):5405–14.DOI: 10.1523/JNEUROSCI.16-17-05405.1996.

31. Bossi, E., et al., Role of anion-cation interactions on the pre-steady-state currents of the rat Na(+)-Cl(-)-dependent GABA cotransporter rGAT1. J Physiol, 2002. 541(Pt 2): p. 343–50.DOI: 10.1113/jphysiol.2001.013457.

32. Kanner, B.I., Transmembrane domain I of the gamma-aminobutyric acid transporter GAT-1 plays a crucial role in the transition between cation leak and transport modes. J Biol Chem, 2003. 278(6): p. 3705–12.DOI: 10.1074/jbc.M210525200.

33. Lu, C.C. and D.W. Hilgemann, GAT1 (GABA:Na+:Cl-) cotransport function. Steady state studies in giant Xenopus oocyte membrane patches. J Gen Physio, 1999. September 114(3):429–44.DOI: 10.1085/jgp.114.3.429

34. Lazzarin, E., et al., Interaction of GAT1 with sodium ions: from efficient recruitment to stabilisation of substrate and conformation. eLife, 2024. 13(RP93271).DOI: 10.7554/elife.93271.1.

35. Motiwala, Z., et al., Structural basis of GABA reuptake inhibition. Nature, 2022.DOI: 10.1038/s41586-022-04814-x.

36. Nayak, S.R., et al., Cryo-EM structure of GABA transporter 1 reveals substrate recognition and transport mechanism. Nat Struct Mol Biol, 2023. 30(7): p. 1023–1032.DOI: 10.1038/s41594-023-01011-w.

37. Fesce, R., et al., The relation between charge movement and transport-associated currents in the rat GABA cotransporter rGAT1. J Physiol, 2002. 545(3): p. 739–50.DOI: 10.1113/jphysiol.2002.026823.

38. Vacca, F., et al., Functional characterization of Atlantic salmon (Salmo salar L.) PepT2 transporters. J Physiol, 2022. 600(10): p. 2377–2400.DOI: 10.1113/JP282781.

39. Mollenhauer HH, Morré DJ, and R. LD, Alteration of intracellular traffic by monensin; mechanism, specificity and relationship to toxicity. Biochim Biophys Acta., 1990.DOI: 10.1016/0304-4157(90)90008-z.

40. Sitte, H.H., E. Singer, A., and P. Scholze, Bi-directional transport of GABA in human embryonic kidney (HEK-293) cells stably expressing the rat GABA transporter GAT-1. British Journal of Pharmacology, 2002. Jan(135(1)): p. 93–102.DOI: 10.1038/sj.bjp.0704446.

41. Varadi, M., et al., AlphaFold Protein Structure Database: massively expanding the structural coverage of protein-sequence space with high-accuracy models. Nucleic Acids Res, 2022. 50(D1): p. D439–D444.DOI: 10.1093/nar/gkab1061.

42. Jumper, J., et al., Highly accurate protein structure prediction with AlphaFold. Nature, 2021. 596(7873): p. 583–589.DOI: 10.1038/s41586-021-03819-2.

43. Coleman, J.A., E.M. Green, and E. Gouaux, X-ray structures and mechanism of the human serotonin transporter. Nature, 2016. 532(7599): p. 334–9.DOI: 10.1038/nature17629.

44. Zhu, A., et al., Molecular basis for substrate recognition and transport of human GABA transporter GAT1. Nat Struct Mol Biol, 2023. 30(7): p. 1012–1022.DOI: 10.1038/s41594-023-00983-z.

45. Mukherjee, S., Betaine and nonalcoholic steatohepatitis: back to the future? World J Gastroenterol, 2011. 17(32): p. 3663–4.DOI: 10.3748/wjg.v17.i32.3663.

46. Van Every, D.W., et al., Betaine Supplementation: A Critical Review of Its Efficacy for Improving Muscle Strength, Power, and Body Composition. Strength & Conditioning Journal, 2021. 43(4): p. 53–61.DOI: 10.1519/ssc.0000000000000622.

47. Chern, M.K. and R. Pietruszko, Evidence for mitochondrial localization of betaine aldehyde dehydrogenase in rat liver: purification, characterization, and comparison with human cytoplasmic E3 isozyme. Biochem Cell Biol., 1999. 77(3): p. 179–87.DOI: 10.1139/o99-030.

48. Slow, S., et al., Plasma Dependent and Independent Accumulation of Betaine in Male and Female Rat Tissues. 2009.DOI: 10.33549/physiolres.931569.

49. Awwad, H.M., et al., Measurement of concentrations of whole blood levels of choline, betaine, and dimethylglycine and their relations to plasma levels. J Chromatogr B Analyt Technol Biomed Life Sci, 2014. 957: p. 41–5.DOI: 10.1016/j.jchromb.2014.02.030.

50. Knight, L.S., et al., Betaine in the Brain: Characterization of Betaine Uptake, its Influence on Other Osmolytes and its Potential Role in Neuroprotection from Osmotic Stress. Neurochem Res, 2017. 42(12): p. 3490–3503.DOI: 10.1007/s11064-017-2397-3.

51. Matthews, A., et al., An indirect response model of homocysteine suppression by betaine: optimising the dosage regimen of betaine in homocystinuria. Br J Clin Pharmacol, 2002. 54(2): p. 140–6.DOI: 10.1046/j.1365-2125.2002.01620.x.

52. Schwahn, B.C., et al., Pharmacokinetics of oral betaine in healthy subjects and patients with homocystinuria. Br J Clin Pharmacol, 2003. 55(1): p. 6–13.DOI: 10.1046/j.1365-2125.2003.01717.x.

53. Wehner, F., et al., Cell volume regulation: osmolytes, osmolyte transport, and signal transduction. Rev Physiol Biochem Pharmacol, 2003. 148: p. 1–80.DOI: 10.1007/s10254-003-0009-x.

54. Schwahn, B.C., et al., Betaine rescue of an animal model with methylenetetrahydrofolate reductase deficiency. Biochem J., 2004.DOI: 10.1042/BJ20040822.

55. Forlani, G., et al., Mutation K448E in the external loop 5 of rat GABA transporter rGAT1 induces pH sensitivity and alters substrate interactions. J Physiol, 2001. 536(Pt 2): p. 479–94.DOI: 10.1111/j.1469-7793.2001.0479c.xd.

56. Clausen, R.P., et al., Selective inhibitors of GABA uptake: synthesis and molecular pharmacology of 4-N-methylamino-4,5,6,7-tetrahydrobenzo[d]isoxazol- 3-ol analogues. Bioorg Med Chem, 2005. 13(3): p. 895–908.DOI: 10.1016/j.bmc.2004.10.029.

57. Jorgensen, L., et al., Structure-Activity Relationship, Pharmacological Characterization, and Molecular Modeling of Noncompetitive Inhibitors of the Betaine/gamma-Aminobutyric Acid Transporter 1 (BGT1). J Med Chem, 2017. 60(21): p. 8834–8846.DOI: 10.1021/acs.jmedchem.7b00924.

58. Latka, K., J. Jonczyk, and M. Bajda, gamma-Aminobutyric acid transporters as relevant biological target: Their function, structure, inhibitors and role in the therapy of different diseases. Int J Biol Macromol, 2020.DOI: 10.1016/j.ijbiomac.2020.04.126.

59. Shahsavar, A. and P. Wellendorph, GABA transport cycle: beyond a GAT feeling. Nat Struct Mol Biol, 2023. 30(7): p. 863–865.DOI: 10.1038/s41594-023-01032-5.

60. Wang, R., et al., Fluorination and Betaine Modification Augment the Blood-Brain Barrier-Crossing Ability of Cylindrical Polymer Brushes. Angew Chem Int Ed Engl, 2022. 61(19): p. e202201390.DOI: 10.1002/anie.202201390.

61. Veszelka, S., et al., Comparison of a Rat Primary Cell-Based Blood-Brain Barrier Model With Epithelial and Brain Endothelial Cell Lines: Gene Expression and Drug Transport. Front Mol Neurosci, 2018. 11: p. 166.DOI: 10.3389/fnmol.2018.00166.

62. Zhang, W., et al., The Role of the GABAergic System in Diseases of the Central Nervous System. Neuroscience, 2021. 470: p. 88–99.DOI: 10.1016/j.neuroscience.2021.06.037.

63. Knight, L.S. and T.A. Knight, Making the case for prophylactic use of betaine to promote brain health in young (15-24 year old) athletes at risk for concussion. Front Neurosci, 2023. 17: p. 1214976.DOI: 10.3389/fnins.2023.1214976.

64. Mayer, F.P., et al., Phase I metabolites of mephedrone display biological activity as substrates at monoamine transporters. Br J Pharmacol, 2016. 173(17): p. 2657–68.DOI: 10.1111/bph.13547.

65. Brüggemann, A., et al., Potassium Channels. Chapter 13: Planar Patch Clamp: Advances in Electrophysiolog. Methods in Molecular Biology, 2009.DOI: 10.1007/978-1-59745-526-8.

66. Rudin, D., et al., (2-Aminopropyl)benzo[beta]thiophenes (APBTs) are novel monoamine transporter ligands that lack stimulant effects but display psychedelic-like activity in mice. Neuropsychopharmacology, 2022. 47(4): p. 914–923.DOI: 10.1038/s41386-021-01221-0.

67. Nadal-Gratacos, N., et al., Structure-Activity Relationship of Novel Second-Generation Synthetic Cathinones: Mechanism of Action, Locomotion, Reward, and Immediate-Early Genes. Front Pharmacol, 2021. 12: p. 749429.DOI: 10.3389/fphar.2021.749429.

68. Monticelli, L., et al., The MARTINI Coarse-Grained Force Field: Extension to Proteins. J Chem Theory Comput, 2008.DOI: 10.1021/ct700324x. PMID: 26621095.

69. de Jong, D.H., et al., Improved Parameters for the Martini Coarse-Grained Protein Force Field. J Chem Theory Comput, 2013. 9(1): p. 687–97.DOI: 10.1021/ct300646g.

70. Wassenaar, T.A., et al., Computational Lipidomics with insane: A Versatile Tool for Generating Custom Membranes for Molecular Simulations. J Chem Theory Comput, 2015. 11(5): p. 2144–55.DOI: 10.1021/acs.jctc.5b00209.

71. van Meer, G., Lipids of the Golgi membrane. Trends Cell Biol., 1998. Jan(8 (1)): p. 29–33.DOI: 10.1016/s0962-8924(97)01196-3.

72. Wassenaar, T.A., et al., Going Backward: A Flexible Geometric Approach to Reverse Transformation from Coarse Grained to Atomistic Models. J Chem Theory Comput, 2014. 10(2): p. 676–90.DOI: 10.1021/ct400617g.

73. Wolf, M.G., et al., Corrigendum: g_membed: Efficient insertion of a membrane protein into an equilibrated lipid bilayer with minimal perturbation. J Comput Chem, 2016. 37(21): p. 2038.DOI: 10.1002/jcc.24386.

74. Szollosi, D. and T. Stockner, Sodium Binding Stabilizes the Outward-Open State of SERT by Limiting Bundle Domain Motions. Cells, 2022. 11(2).DOI: 10.3390/cells11020255.

75. Lindorff-Larsen, K., et al., Improved side-chain torsion potentials for the Amber ff99SB protein force field. Proteins, 2010. 78(8): p. 1950–8.DOI: 10.1002/prot.22711.

76. Jambeck, J.P. and A.P. Lyubartsev, Another Piece of the Membrane Puzzle: Extending Slipids Further. J Chem Theory Comput, 2013. 9(1): p. 774–84.DOI: 10.1021/ct300777p.

77. Abraham, M.J., et al., GROMACS: High performance molecular simulations through multi-level parallelism from laptops to supercomputers. SoftwareX, 2015. 1-2: p. 19–25.DOI: 10.1016/j.softx.2015.06.001.

78. Sohail, A., et al., The Environment Shapes the Inner Vestibule of LeuT. PLoS Comput Biol, 2016. 12(11): p. e1005197.DOI: 10.1371/journal.pcbi.1005197.

79. Goda, K., et al., Human ABCB1 with an ABCB11-like degenerate nucleotide binding site maintains transport activity by avoiding nucleotide occlusion. PLoS Genet, 2020. 16(10): p. e1009016.DOI: 10.1371/journal.pgen.1009016.

80. Bussi, G., D. Donadio, and M. Parrinello, Canonical sampling through velocity rescaling. J Chem Phys, 2007. 126(1): p. 014101.DOI: 10.1063/1.2408420.

81. Parrinello, M. and A. Rahman, Polymorphic transitions in single crystals: A new molecular dynamics method. J. Appl. Phys., 1981. December.DOI: 10.1063/1.328693.

82. Darden, T., D. York, and L. Pedersen, Particle mesh Ewald: An N⋅log(N) method for Ewald sums in large systems. The Journal of Chemical Physics, 1993. 98(12): p. 10089–10092.DOI: 10.1063/1.464397.

83. Mager, S., et al., Steady states, charge movements, and rates for a cloned GABA transporter expressed in Xenopus oocytes. Neuron., 1993. February 10(2):177–88.DOI: 10.1016/0896-6273(93)90309-f.

