## Supplementary data for "Unveiling the crucial role of betaine: Modulation of GABA homeostasis via SLC6A1 transporter (GAT1)"

**Results**

Glycine does not induce transport currents in GAT1, while betaine-induced currents can be blocked by GAT1 inhibitors.

Apart from being an important inhibitory neurotransmitter, glycine is also an osmolyte and a precursor to betaine. We studied the interaction of glycine with GAT1, as a control condition. The *X. laevis* oocytes expressing rGAT1 were clamped at the holding potential of -60mV and perfused with different glycine concentrations (1-50 mM) (Figure S1 A). We did not observe any inward currents due to glycine, but a response like in non-injected control oocytes.

In the main text, we showed that SKF89976a can inhibit betaine-induced transport currents in oocytes expressing rGAT1 (Figure 1). We also studied the effects of other GAT1 inhibitors, tiagabine and NO-711^1^ on the betaine transport by GAT1. The *X. laevis* oocytes expressing rGAT1 were clamped at the holding potential of –60 mV and were perfused with GABA 100µM and betaine 10mM with and without tiagabine 3 µM and NO-711 10 µM (Figure S1 B). The large inward transport current induced by GABA 100 µM was reduced strongly by both tiagabine 3 µM and NO-711 10 µM. Similarly, the currents induced by betaine 10 mM were blocked by both tiagabine 3 µM and NO-711 10 µM. Hence, like GABA, betaine transport by GAT1 can be inhibited by the GAT1 blockers tiagabine and NO-711.

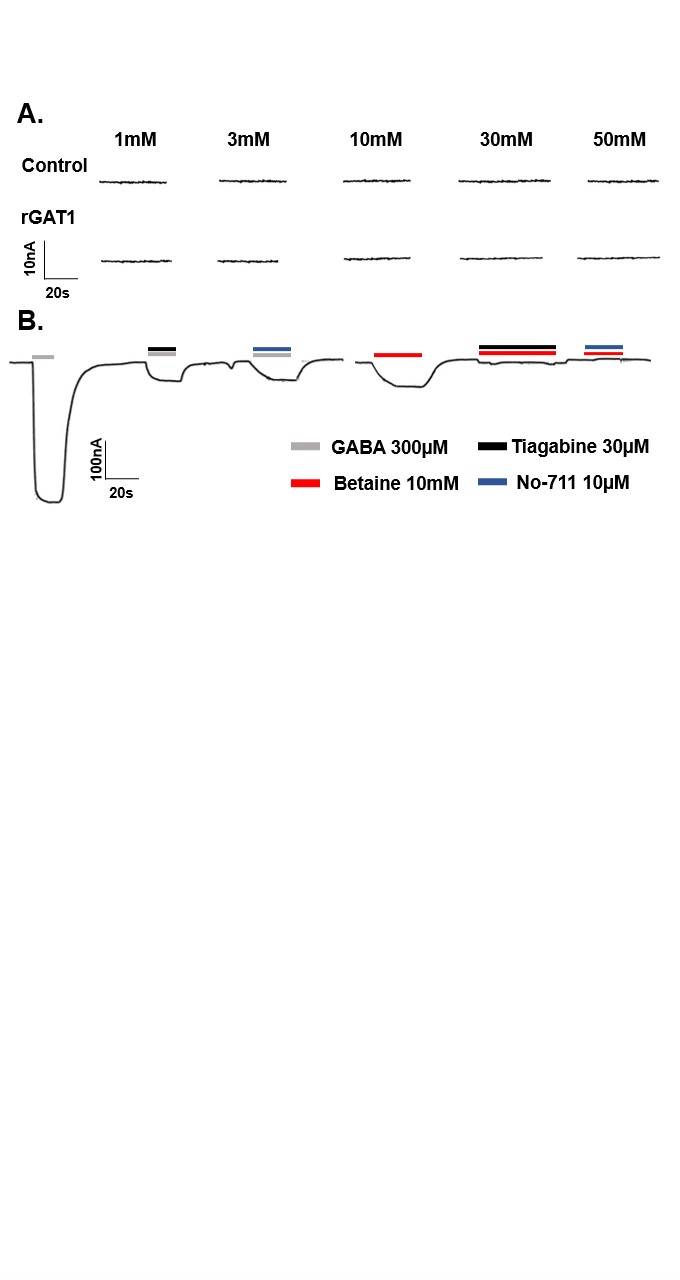
***Figure S1: Interaction of glycine with rGAT1 and GAT1 blockers with betaine transport by rGAT1.* A.** The representative current traces of the perfusion of different glycine concentrations (as labelled) on the non-injected oocyte and the oocyte expressing rGAT1, at holding potential –60 mV. **B.** The representative trace of *X. laevis* oocyte expressing rGAT1, perfused with GABA 100 µM and betaine 10 mM with and without tiagabine 3 µM and NO-711 10 µM, at holding potential –60 mV. The inward currents induced by both GABA and betaine were blocked by both tiagabine and NO-711.

#### Automated patch-clamping also shows that betaine can be translocated by GAT1.

Perfusion of betaine in the presence of 140 mM NaCl was able to generate appreciable inward steady-state currents on CHO cells overexpressing hGAT1 starting from 10 mM substrate (Figure S2 A), at -80 mV. The maximum current amplitude (around -40 pA) was reached upon the addition of 300 mM betaine to the external solution. The addition of 600 mM betaine generated spikes, in some cases, at around 7-8 s after addition (Figure S1 A), probably due to heavy osmolar stress. Applying betaine in a rising concentration sequence (3-600 mM) generated inward-directed currents proportionally increasing with betaine concentrations, and the data was fitted using Hill’s model (Figure S2 B). The EC_50_ for betaine on hGAT1 was calculated to be 27.00 ± 3.24 mM with a Hill’s coefficient of 1.57 ± 0.28, at the holding potential of -80 mV and room temperature. Of the eight wells of the 4X chip used, six were suitable for carrying out the experiment (one showed a seal resistance <300 MΩ and another showed leakage after the Ca^2+^-washing step), therefore n=6.

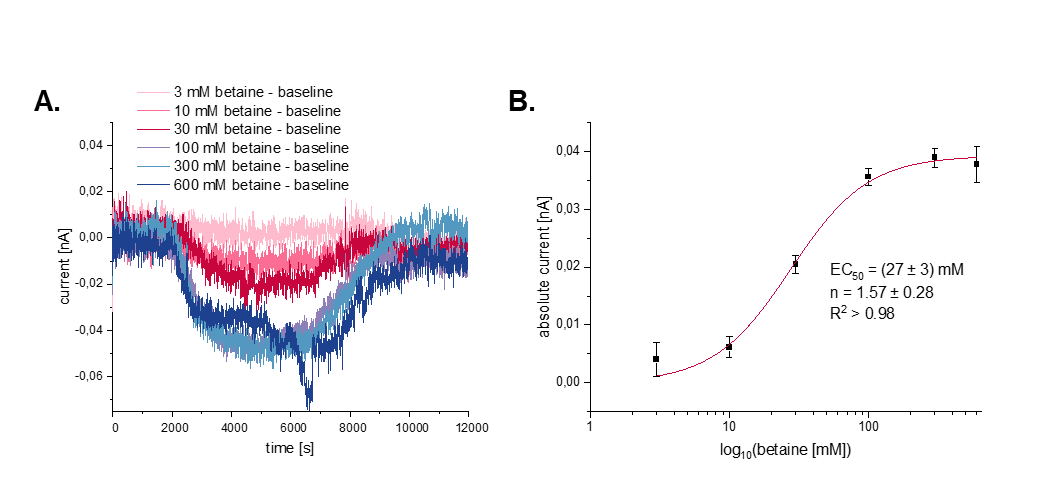

***Figure S2: Dose-response of betaine in CHO cells overexpressing hGAT1, using automated patch clamp.* A.** The representative current traces induced by perfusion of different betaine concentrations (as labelled). The spike around 7 s at 600 mM betaine is probably caused by the osmolar stress. **B.** The kinetic analysis of betaine induced currents shows EC_50_ = 27.00 ± 3.24 mM and Hill’s coefficient as 1.57 ± 0.28. Data are shown as the mean ± SD from n=6 and fitted using the Hill’s model. The holding potential for all measurements was –80 mV.

### The pre-steady state analysis of the voltage-step response for betaine in *X. laevis* oocytes expressing rGAT1.

As shown in the main text, betaine is a slower substrate of rGAT1. The voltage-step experiment of the *X. laevis* oocytes expressing rGAT1 was performed for different betaine concentrations (0.1-50 mM). Here we show the analysis for the betaine concentrations that were not shown in the main text (Figure 2). Just like other betaine concentrations, we see an increase in voltage-dependent inward transport current with increase in extracellular betaine (Figure S3 A). Similarly, the total charge dislocation (Q) and the decay time constant (τ) decreased with the increase in betaine concentration (Figure S3 B, C). Also, the unidirectional rate constants outrate (α) and inrate (β) increased with the increase in betaine concentration, as with higher concentration more betaine and Na^+^ ions can be translocated due to the chemical gradient, resulting in the reduction of the pre-steady state components (Figure S3 D).

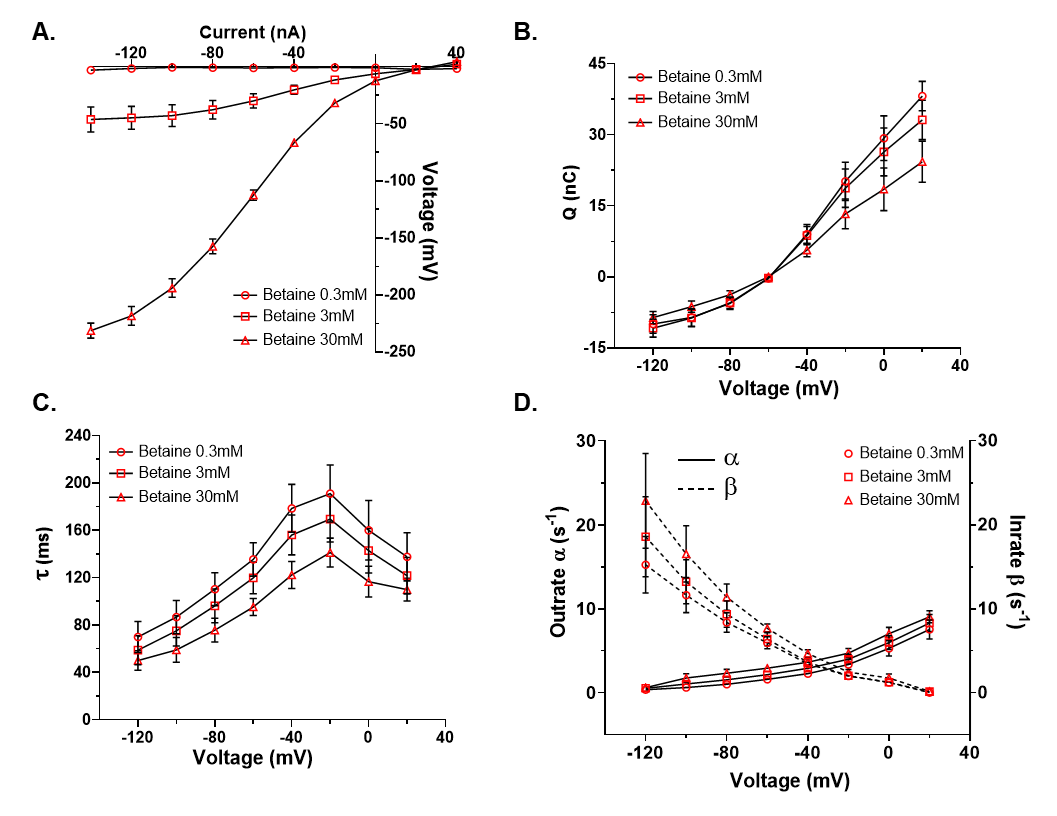
***Figure S3: The pre-steady state analysis of betaine transport by rGAT1.*** The current response for each condition was collected by giving 0.8 s long squared pulse at –20 mV of voltage jump. **A.** The current and voltage (I-V) relationship from –140 mV to +40 mV. **B.** The total charge dislocation and voltage (Q-V) relationship. **C.** The decay time constant and voltage (τ-V) relationship. **D.** The relationships of unidirectional rate constants outrate (α) and inrate (β, shown as dashed line) with voltage. All the reported values were collected in the presence of ND98 alone and/or with betaine 0.3, 3, 30 mM. In B-D the voltage tested was from –120 mV to +20 mV. All values are shown as mean ± SEM of 3/1 n/N at the holding potential -60 mV.

The standard calibration curve for GABA and betaine detection in *X. laevis* oocytes expressing rGAT1

To verify the sensitivity and linearity of the LC-MS/MS response to GABA and betaine their standard solutions were created, and calibration curves were obtained. The different concentrations of GABA and betaine (2, 4, 8, 16 µM) were diluted in the LC-MS grade H_2_O, with 0.1% formic acid. Calibration curves showing the relative GABA and betaine peak areas plotted against analytes concentrations (range 2–16 µM) were created (Figure S4).

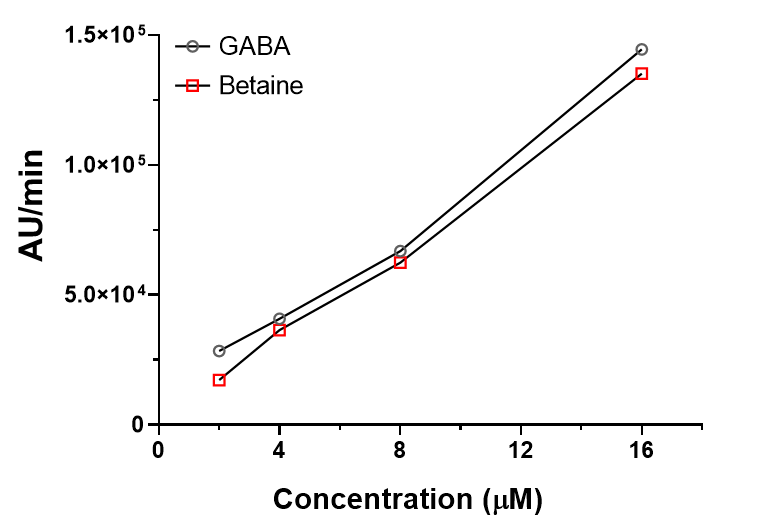

***Figure S4: The standard calibration curve for the detection of GABA and betaine in X. laevis oocytes using LC-MS/MS.*** The analysis of different GABA and betaine concentrations (2, 4, 8, 16 µM) diluted with LC-MS grade H_2_O, with 0.1% formic acid shows a linear increase in the presence of GABA and betaine. All values are shown as the arbitrary unit per oocyte per minute from 5/1 n/N.

#### Molecular docking of GABA in hGAT1 in the outward-open state

We performed molecular docking of GABA in the Alphafold model of hGAT1 in outward-open state with two Na^+^ and a Cl^‑^. The docking simulations gave a fitness score of 50.52, and showed that GABA bound with Na^+^ at Na1 site, formed short-range (d < 3 Å) contacts with Y60, G65, Y140, S396, and T400, and medium-range (3 Å < d < 5 Å) contacts with L64, F294, S295 (Figure S5 A). These data match with the existing literature and the information obtained from the recent cryo-EM structures, and were used as the positive control for the docking experiments of betaine and hGAT1 (Figure S5 B)^2-5^.

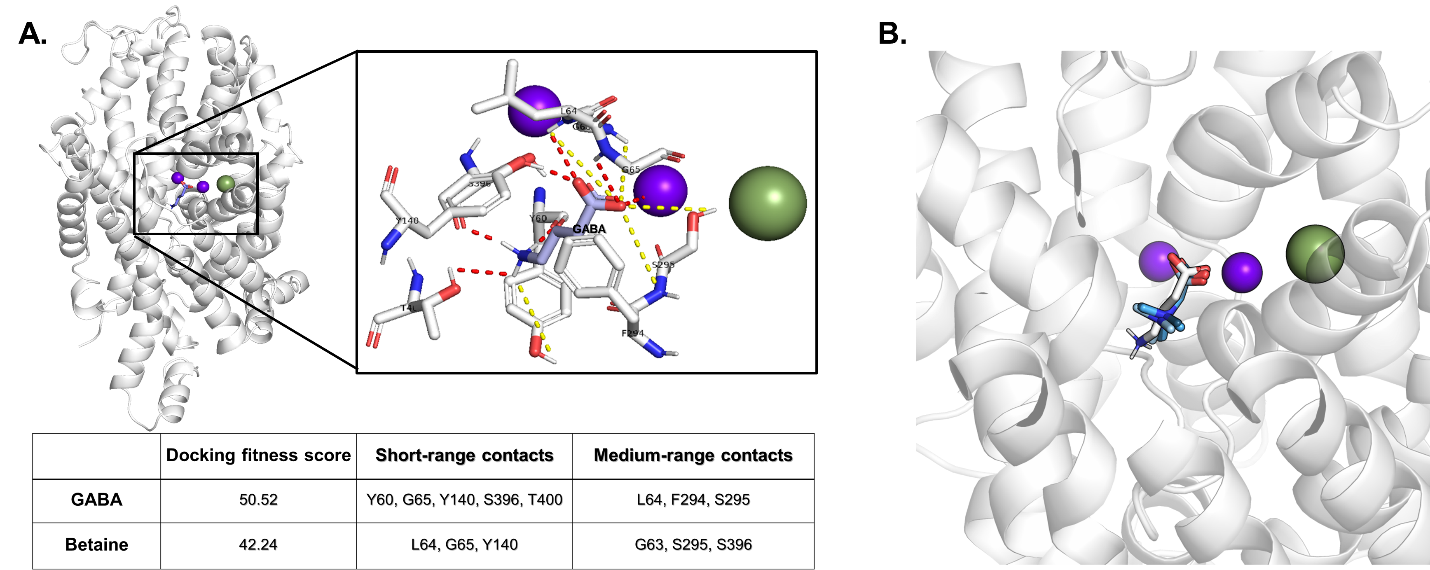

***Figure S5: Molecular docking of GABA in Alphafold hGAT1 model in outward-open state.* A.** The zoomed-in view of GABA docked in outward-open hGAT1 with two sodium ions and a chloride ion. GABA formed short-range contacts (d < 3 Å) with five residues and medium-range contacts (3 Å < d < 5 Å) with three. The table below summarizes the docking fitness score, and the residues involved in GABA binding. **B.** The docking of betaine with outward-open hGAT1 with GABA as a reference shown in three docking poses with the best docking fitness score.

#### Betaine does not show a selective inhibition of GABA uptake by *X. laevis* oocytes heterologously expressing cBGT-1.

The *X. laevis* oocytes expressing cBGT-1 were clamped at the holding potential of –60 mV, to perform GABA-betaine competitive assay experiments. The half maximal concentration for both GABA and betaine were calculated and found to be K_0.5,GABA_~30 µM and K_0.5,betaine_~1.7 mM (as shown in the table of Figure 1). To investigate the GABA-betaine relationship, the oocytes were perfused with low betaine concentration of 0.3 mM with GABA 30 µM and 300 µM, and high betaine concentration of betaine 3 mM with the same GABA concentrations (Figure S6 A). The heatmap analysis of the current recorded for these conditions showed the absence of any inhibitory effects of betaine on GABA induced transport current, neither at low nor at high concentrations (Figure S6 B). The peculiar inhibitory effects of betaine on GABA seem to be particular in the case of GAT1.

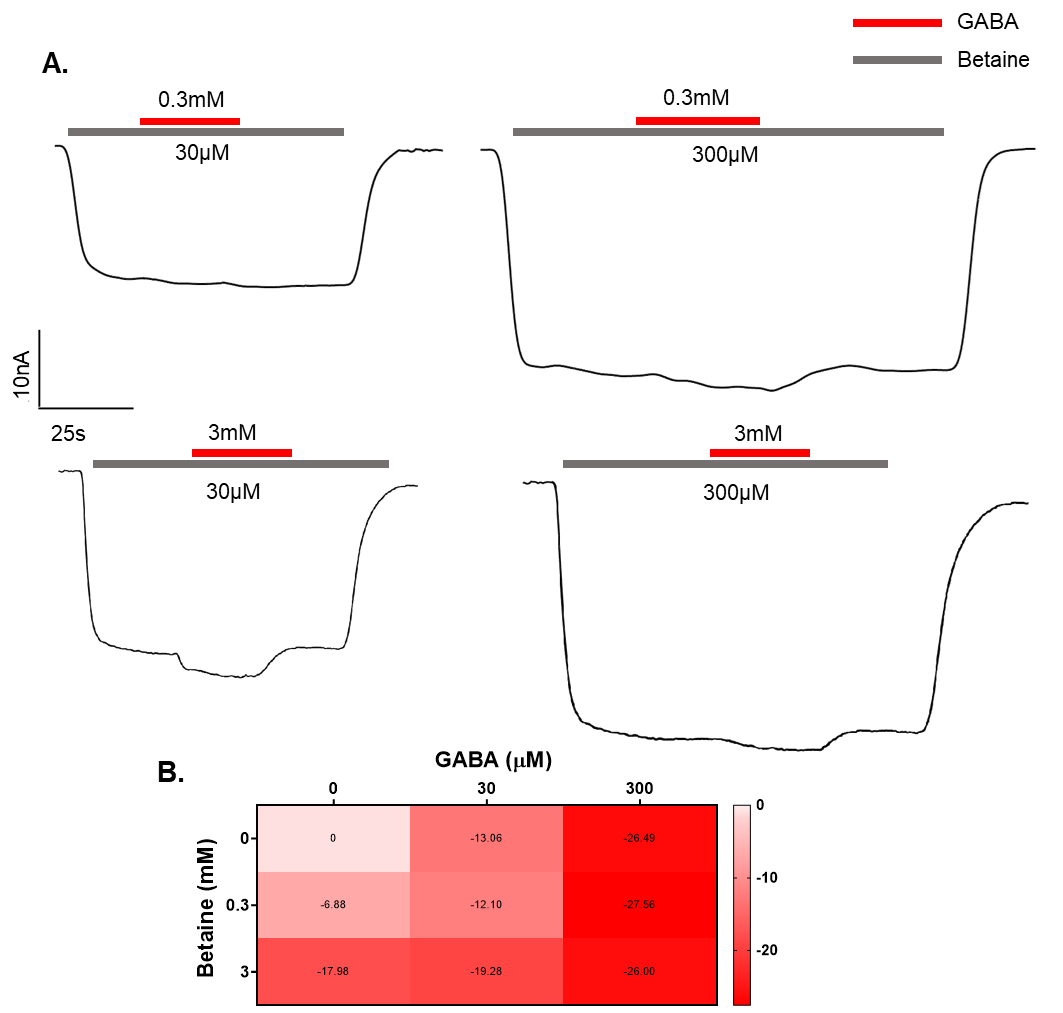

***Figure S6: Betaine does not selectively inhibit GABA transport in X. laevis oocytes expressing cBGT-1.* A.** The representative traces of the current induced by betaine 0.3 and 3 mM with GABA 30 and 300 µM. **B.** The heatmap analysis of the transport current recorded for betaine 0.3 and 3 mM with and without GABA 30 and 300 µM, shows no inhibitory effects induced by betaine. Data are shown as the mean ± SEM of 3/2. The holding potential for all measurements was –60 mV.

#### In GAT1, the selective inhibition by betaine is specific to GABA and absent with its secondary substrate nipecotic acid.

The cyclic analogue of GABA i.e. nipecotic acid has been known for almost five decades as the competitive inhibitor of GAT1^6^^,^^7^. There were further chemical modulations done on nipecotic acid to derive potent GAT1 inhibitors that do not act like a substrate and can cross blood-brain barrier e.g. tiagabine, SKF89976a etc. We have already shown that these blockers inhibit the betaine transport by GAT1 (Figure 1). Since nipecotic acid is also a substrate of GAT1, we performed the competitive assay of betaine and nipecotic assay on *X. laevis* oocytes expressing rGAT1 and compared the results with the competitive assay of GABA and nipecotic acid. Given that the half-maximal concentration for nipecotic acid in GAT1 is ~20 µM ^7^, we performed competition of nipecotic acid 20 µM and 200 µM (saturating concentration) with betaine 100 µM (where maximum GABA inhibition was observed, Figure 6) and GABA 10 µM (~K_0.5_) (Figure S7 A). We observed no inhibition of the transport current for GABA 10 µM by Nipecotic acid 20 µM, whereas at high concentration of nipecotic acid (200 µM) the overall transport current was higher (-81.41 ± 5.87 nA) than GABA 10µM (-57.99 ± 2.91 nA) or nipecotic acid 200 µM (-70.07 ± 2.12 nA) alone (Figure S8 B). Whereas with betaine 100 µM, no such changes were observed either with nipecotic acid 20 µM or 200 µM (Figure S7 B). Hence, the peculiar GABA inhibition induced by low betaine concentrations in GAT1 seems to be GABA specific.

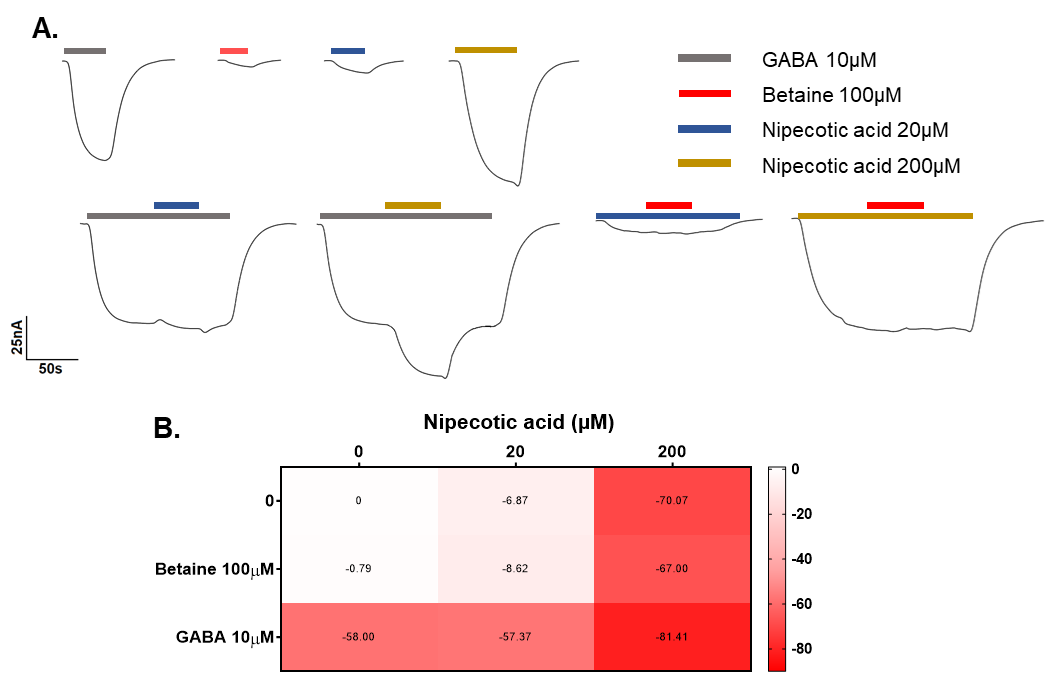
***Figure S7: The competitive assay of nipecotic acid with GABA and betaine in X. laevis oocytes expressing rGAT1.* A.** The representative traces of the current recorded with nipecotic acid 20 and 200 µM alone, and with GABA 10 µM and betaine 100 µM. **B.** The heatmap analysis of nipecotic acid 20 and 200 µM with GABA 10 µM and betaine 100 µM that shows no alteration in transport current recorded with nipecotic acid in the presence of betaine, unlike GABA. Data are shown as the mean ± SEM of 6/1. The holding potential for all measurements was –60 mV.

|  | **ND98** | **Betaine 0.1mM** | **GABA 10µM** | **GABA 10µM+ betaine 0.1mM** |
| --- | --- | --- | --- | --- |
| **τ_max_ (ms)** | 191.72±18.3 | 192.67±18.5 | 137.01±14.2 | 187.83±30.8 |
| **Q_max_ (nC)** | 38.14±1.0 | 43.59±2.3 | 25.74±3.3 | 30.29±2.1 |
| **V_0.5_ (mV)** | -32.86±1.1 | -27.16±2.6 | -43.19±3.4 | -39.18±2.5 |
| **σ (mV)** | 20.36±0.7 | 23.49±1.6 | 25.21±3.5 | 21.81±1.3 |

***Table S1:*** The maximum value of decay time constant (τ_max_) and the fitting parameters from Q-V relationship in rGAT1 obtained using the Boltzmann distribution equation. Q_max_ is the maximum moveable charge, V_0.5_ is the voltage where half of the charge can be moved, and σ is the slope factor of the sigmoidal curve. All data are means ± SEM of 7/3 n/N.

|  | **Inrate β (s^-1^)** | | **Outrate α (s^-1^)** | |
| --- | --- | --- | --- | --- |
| **Voltage (mV)** | **GABA 10µM** | **GABA 10µM+ betaine 0.1mM** | **GABA 10µM** | **GABA 10µM+ betaine 0.1mM** |
| **-120** | 17.78±5.9 | 13.11±3.3 | -0.30±0.2 | -0.16±0.1 |
| **-100** | 12.33±3.1 | 10.12±2.4 | 1.22±0.4 | 0.61±0.1 |
| **-80** | 8.12±1.5 | 7.34±1.5 | 2.61±0.6 | 1.21±0.1 |
| **-60** | 4.82±1.1 | 4.55±0.8 | 3.62±0.9 | 1.86±0.1 |
| **-40** | 2.28±1.5 | 2.56±0.4 | 5.18±0.7 | 3.03±0.4 |
| **-20** | 0.66±0.1 | 1.17±0.1 | 7.08±0.8 | 4.40±0.5 |
| **0** | 0.31±0.5 | 0.64±0.1 | 10.32±0.7 | 6.70±0.6 |
| **20** | -0.41±0.5 | 0.01±0.1 | 12.63±0.6 | 8.84±0.7 |

***Table S2:*** Unidirectional rate constants outrate α (s^-1^) and inrate β (s^-1^) for voltages -120 mV to +20 mV. All data are means ± SEM of 7/3 n/N.

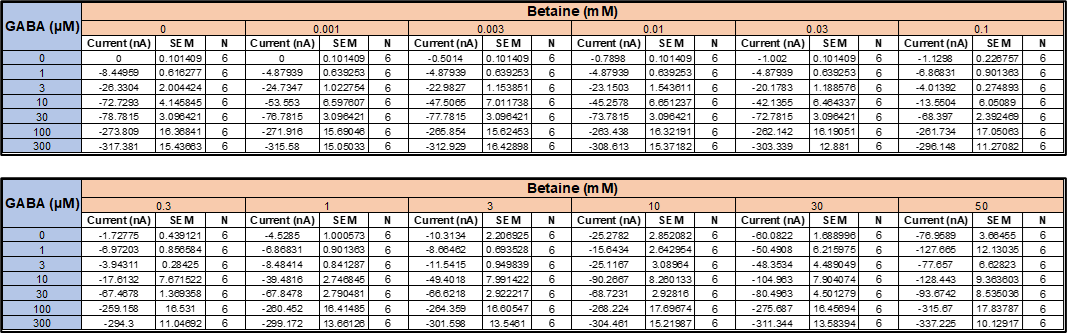

***Table S3:*** The transport current values for GABA-betaine competition in *X. laevis* oocytes expressing rGAT1, displayed as a heatmap in Figure 6 B. Data are shown as mean ± SEM of 6/2 n/N.

Code S1: Production parameters for the molecular dynamic simulation of betaine with hGAT1.

VARIOUS PREPROCESSING OPTIONS

Preprocessor information: use cpp syntax.

e.g.: -I/home/joe/doe -I/home/mary/roe

include	=

e.g.: -DPOSRES -DFLEXIBLE (note these variable names are case sensitive)

define	= -DPOSRES

RUN CONTROL PARAMETERS

integrator	= md

Start time and timestep in ps

tinit	= 0

dt	= 0.002

nsteps	= 500000000 1000 ns

For exact run continuation or redoing part of a run

init_step	= 0

Part index is updated automatically on checkpointing (keeps files separate)

simulation_part	= 1

mode for center of mass motion removal

comm-mode	= linear

number of steps for center of mass motion removal

nstcomm	= 100

group(s) for center of mass motion removal

comm-grps	= ProtLigIon membrane Water_and_ions

LANGEVIN DYNAMICS OPTIONS

Friction coefficient (amu/ps) and random seed

bd-fric	= 0

ld-seed	= 1993

ENERGY MINIMIZATION OPTIONS

Force tolerance and initial step-size

emtol	= 1000

emstep	= 0.0001

Max number of iterations in relax-shells

niter	= 20

Step size (ps^2) for minimization of flexible constraints

fcstep	= 0.001

Frequency of steepest descents steps when doing CG

nstcgsteep	= 50

nbfgscorr	= 100

TEST PARTICLE INSERTION OPTIONS

rtpi	= 0.05

OUTPUT CONTROL OPTIONS

Output frequency for coords (x), velocities (v) and forces (f)

nstxout	= 5000 10 ps

nstvout	=

nstfout	= 0

Output frequency for energies to log file and energy file

nstlog	= 1000

nstcalcenergy	= 100

nstenergy	= 1000

Output frequency and precision for .xtc file

nstxout-compressed	= 0

compressed-x-precision	= 1000

This selects the subset of atoms for the compressed trajectory file. You can select multiple groups. By default, all atoms will be written.

compressed-x-grps	=

Selection of energy groups

energygrps	=

NEIGHBORSEARCHING PARAMETERS

cut-off scheme (Verlet: particle-based cut-offs, group: using charge groups)

cutoff-scheme	= Verlet

nblist update frequency

nstlist	= 50

ns algorithm (simple or grid)

ns-type	= Grid

Periodic boundary conditions: xyz, no, xy

pbc	= xyz

periodic_molecules	= no

Allowed energy error due to the Verlet buffer in kJ/mol/ps per atom,

a value of -1 means: use rlist

verlet-buffer-tolerance = 0.005

nblist cut-off

rlist	= 0.9

long-range cut-off for switched potentials

rlistlong	= -1

nstcalclr	= -1

OPTIONS FOR ELECTROSTATICS AND VDW

Method for doing electrostatics

coulombtype	= PME

coulomb-modifier	= Potential-shift-Verlet

rcoulomb-switch	=

rcoulomb	= 0.9

Relative dielectric constant for the medium and the reaction field

epsilon_r	= 1.0

epsilon_rf	= 1

Method for doing Van der Waals

vdw-type	= Cut-off

vdw-modifier	= Potential-shift-Verlet

cut-off lengths

rvdw-switch	=

rvdw	= 0.9

Apply long range dispersion corrections for Energy and Pressure

DispCorr	= EnerPres

Extension of the potential lookup tables beyond the cut-off

table-extension	= 1

Separate tables between energy group pairs

energygrp-table	=

Spacing for the PME/PPPM FFT grid

fourierspacing	= 0.12

FFT grid size, when a value is 0 fourierspacing will be used

fourier_nx	= 0

fourier_ny	= 0

fourier_nz	= 0

EWALD/PME/PPPM parameters

pme_order	= 4

ewald_rtol	= 1e-05

ewald-rtol-lj	= 0.001

lj-pme-comb-rule	= Geometric

ewald_geometry	= 3d

epsilon_surface	= 0

IMPLICIT SOLVENT ALGORITHM

implicit_solvent	= No

GENERALIZED BORN ELECTROSTATICS

Algorithm for calculating Born radii

gb-algorithm	= Still

Frequency of calculating the Born radii inside rlist

nstgbradii	= 1

Cutoff for Born radii calculation the contribution from atoms between rlist and rgbradii is updated every nstlist steps

rgbradii	= 1

Dielectric coefficient of the implicit solvent

gb-epsilon-solvent	= 80

Salt concentration in M for Generalized Born models

gb-saltconc	= 0

Scaling factors used in the OBC GB model. Default values are OBC(II)

gb-obc-alpha	= 1

gb-obc-beta	= 0.8

gb-obc-gamma	= 4.85

gb-dielectric-offset	= 0.009

sa-algorithm	= Ace-approximation

Surface tension (kJ/mol/nm^2) for the SA (nonpolar surface) part of GBSA

The value -1 will set default value for Still/HCT/OBC GB-models.

sa-surface-tension	= -1

OPTIONS FOR WEAK COUPLING ALGORITHMS

Temperature coupling

tcoupl	= v-rescale

nsttcouple	= -1

nh-chain-length	= 10

print-nose-hoover-chain-variables = no

Groups to couple separately

tc-grps	= ProtLigIon membrane Water_and_ions

Time constant (ps) and reference temperature (K)

tau-t	= 0.5	0.5	0.5

ref-t	= 310	310	310

pressure coupling

Pcoupl	= Parrinello-Rahman Berendsen for EM and equilibrationParrinello-Rahman for production

Pcoupltype	= Semiisotropic

nstpcouple	= -1

Time constant (ps), compressibility (1/bar) and reference P (bar)

tau-p	= 20.1	Use 5 for Berendsen 20 for Parrinello-Rahman

compressibility	= 4.5e-05	4.5e-05

ref-p	= 1.0	1.0

Scaling of reference coordinates, No, All or COM

refcoord_scaling	= All

SIMULATED ANNEALING

Type of annealing for each temperature group (no/single/periodic)

annealing	= no

Number of time points to use for specifying annealing in each group

annealing-npoints	=

List of times at the annealing points for each group

annealing-time	=

Temp. at each annealing point, for each group.

annealing-temp	=

GENERATE VELOCITIES FOR STARTUP RUN

gen-vel	= no

gen-temp	= 310.0

gen-seed	= -1

OPTIONS FOR BONDS

constraints	= h-bonds

Type of constraint algorithm

constraint-algorithm	= lincs

Do not constrain the start configuration

continuation	= no

Use successive overrelaxation to reduce the number of shake iterations

Shake-SOR	= yes

Relative tolerance of shake

shake-tol	= 0.0001

Highest order in the expansion of the constraint coupling matrix

lincs-order	= 4

Number of iterations in the final step of LINCS. 1 is fine for normal simulations, but use 2 to conserve energy in NVE runs.

For energy minimization with constraints it should be 4 to 8.

lincs-iter	= 2

Lincs will write a warning to the stderr if in one step a bond rotates over more degrees than

lincs-warnangle	= 30

Convert harmonic bonds to morse potentials

morse	= no

ENERGY GROUP EXCLUSIONS

Pairs of energy groups for which all non-bonded interactions are excluded

energygrp-excl	=

WALLS

Number of walls, type, atom types, densities and box-z scale factor for Ewald

nwall	= 0

wall_type	= 9-3

wall_r_linpot	= -1

wall-atomtype	=

wall-density	=

wall_ewald_zfac	= 3

COM PULLING

Pull type: no, umbrella, constraint or constant-force

pull	= no

ENFORCED ROTATION

Enforced rotation: No or Yes

rotation	= no

Group to display and/or manipulate in interactive MD session

IMD-group	=

NMR refinement stuff

Distance restraints type: No, Simple or Ensemble

disre	= No

Force weighting of pairs in one distance restraint: Conservative or Equal

disre-weighting	= Conservative

Use sqrt of the time averaged times the instantaneous violation

disre-mixed	= no

disre-fc	= 100

disre-tau	= 0

Output frequency for pair distances to energy file

nstdisreout	= 5000

Orientation restraints: No or Yes

orire	= no

Orientation restraints force constant and tau for time averaging

orire-fc	= 0

orire-tau	= 0

orire-fitgrp	=

Output frequency for trace(SD) and S to energy file

nstorireout	= 100

Free energy variables

free-energy	= no

couple-moltype	=

couple-lambda0	= vdw-q

couple-lambda1	= vdw-q

couple-intramol	= no

init-lambda	= 0

init-lambda-state	= -1

delta-lambda	= 0

nstdhdl	= 50

fep-lambdas	=

mass-lambdas	=

coul-lambdas	=

vdw-lambdas	=

bonded-lambdas	=

restraint-lambdas	=

temperature-lambdas	=

calc-lambda-neighbors	= 1

init-lambda-weights	=

dhdl-print-energy	= no

sc-alpha	= 0

sc-power	= 1

sc-r-power	= 6

sc-sigma	= 0.3

sc-coul	= no

separate-dhdl-file	= yes

dhdl-derivatives	= yes

dh_hist_size	= 0

dh_hist_spacing	= 0.1

Non-equilibrium MD stuff

acc-grps	=

accelerate	=

freezegrps	=

freezedim	=

cos-acceleration	= 0

deform	=

simulated tempering variables

simulated-tempering	= no

simulated-tempering-scaling = geometric

sim-temp-low	= 300

sim-temp-high	= 300

Electric fields

Format is number of terms (int) and for all terms an amplitude (real)

and a phase angle (real)

E-x	=

Time dependent (pulsed) electric field. Format is omega, time for pulse peak, and sigma (width) for pulse. Sigma = 0 removes pulse, leaving the field to be a cosine function.

E-xt	=

E-y	=

E-yt	=

E-z	=

E-zt	=

Ion/water position swapping for computational electrophysiology setups

Swap positions along direction: no, X, Y, Z

swapcoords	= no

AdResS parameters

adress	= no

User defined thingies

user1-grps	=

user2-grps	=

userint1	= 0

userint2	= 0

userint3	= 0

userint4	= 0

userreal1	= 0

userreal2	= 0

userreal3	= 0

userreal4	= 0

### **References**

1	Clausen, R. P. *et al.* Selective inhibitors of GABA uptake: synthesis and molecular pharmacology of 4-N-methylamino-4,5,6,7-tetrahydrobenzo[d]isoxazol-3-ol analogues. *Bioorg Med Chem* **13**, 895-908 (2005). <https://doi.org:10.1016/j.bmc.2004.10.029>

2	Zhu, A. *et al.* Molecular basis for substrate recognition and transport of human GABA transporter GAT1. *Nat Struct Mol Biol* **30**, 1012-1022 (2023). <https://doi.org:10.1038/s41594-023-00983-z>

3	Motiwala, Z. *et al.* Structural basis of GABA reuptake inhibition. *Nature* (2022). <https://doi.org:10.1038/s41586-022-04814-x>

4	Zafar, S. & Jabeen, I. Structure, Function, and Modulation of gamma-Aminobutyric Acid Transporter 1 (GAT1) in Neurological Disorders: A Pharmacoinformatic Prospective. *Front Chem* **6**, 397 (2018). <https://doi.org:10.3389/fchem.2018.00397>

5	Nayak, S. R. *et al.* Cryo-EM structure of GABA transporter 1 reveals substrate recognition and transport mechanism. *Nat Struct Mol Biol* **30**, 1023-1032 (2023). <https://doi.org:10.1038/s41594-023-01011-w>

6	Johnston, G. A., Krogsgaard-Larsen, P. & Stephanson, A. Betel nut constituents as inhibitors of gamma-aminobutyric acid uptake. *Nature* **December 18; 258(5536):627-8** (1975). <https://doi.org:10.1038/258627a0>

7	Sarup, A., Larsson OM & A., S. GABA transporters and GABA-transaminase as drug targets. *Curr Drug Targets CNS Neurol Disord* **Aug**, 269-277 (2003 ). <https://doi.org:10.2174/1568007033482788>
